# Transition metal-triggered immunity via an *Arabidopsis* NLR pair

**DOI:** 10.64898/2026.04.10.717661

**Authors:** Cheng Gao, Sisi Chen, Jie Chen, Zhong Tang, Xin-Yuan Huang, Peng Wang, Suomeng Dong, Jeffery L. Dangl, Li Wan, Fang-Jie Zhao

**Author notes:** Corresponding authors: Li Wan, Fang-Jie Zhao. These authors contributed equally to this work (Cheng Gao, Sisi Chen).

## Abstract

Plants are challenged by multiple biotic (e.g. pathogens) and abiotic (e.g. heavy metals) stressors. Some transition metals can enhance plant’s defense against pathogens, but the mechanism remains unclear. Here, we demonstrate that an *Arabidopsis* head-to-head gene pair of intracellular nucleotide-binding leucine-rich repeat (NLR) receptors, both expressed in the endodermis of roots, antagonistically control transition metal-triggered immunity. One NLR, STM2 binds transition metal ions, such as Cd^2+^, Cu^2+^ and Zn^2+^, with its LRR domain to enhance its NAD^+^ hydrolytic activity and immune responses via the EDS1/PAD4/ADR1 module, triggering enhanced resistance to bacterial wilt *Ralstonia solanacearum*. The other NLR, STM1 suppresses STM2 to protect plants from transition metal-triggered immunity and growth inhibition in the presence of excess metals. STM1 also dampens resistance to the pathogen. Our study defines an NLR activated by transition metals and reveals a trade-off between susceptibility to pathogens and sensitivity to transition metals that are pervasive in soil.

## Main

Plants face myriad biotic (e.g. pathogens) and abiotic stressors (e.g. excess metal ions) in the environment and, to survive, evolved various resistance mechanisms. Resistance to pathogen infection is provided by two integrated branches of the host innate immunity, pathogen-associated molecular pattern (PAMP)-triggered immunity (PTI) and effector-triggered immunity (ETI) ^1^. Plant NLR proteins function as primarily intracellular immune receptors, activating ETI by recognizing invasive biological effector proteins entering plant cells ^1,2^.

ETI is often associated with localized plant cell death, referred to as the hypersensitive response (HR) ^3^. There are hundreds of *NLR* genes in most angiosperm plant genomes. NLR proteins carry either a TIR domain or a coiled-coil (CC) domain at the N-terminus and thus are named as TIR-NLR (TNL) and CC-NLR (CNL), respectively. A subset of NLR genes are encoded in head-to-head gene pairs, such as *Arabidopsis* TNL pair *RRS1/RPS4* ^4,5^ and rice CNL pair *RGA5/RGA4* ^6^, which function cooperatively as a sensor for pathogen effector proteins and an executor of immunity. Antagonistic NLRs were also reported. For example, PigmR confers disease resistance but reduces grain yield of rice, whereas PigmS, which is not encoded in a head-to-head gene pair with PigmR, inhibits the disease resistance function of PigmR and increases yield, suggesting a cost associated with PigmR expression in the absence of the corresponding pathogen^7^. However, the function of most NLR gene pairs and the effectors they respond to remain unknown. Although plant immunity is essential for disease resistance, it must be tightly controlled to avoid fitness cost as there is often a trade-off between growth and disease resistance. For example, the NLR protein ACQOS/VICTR contributes to bacterial resistance in the absence of osmotic stress, but causes detrimental autoimmunity and reduced osmo-tolerance in the presence of osmotic stress ^8^.

Plants require at least 14 mineral elements, including several transition metals (e.g. Fe, Mn, Zn, Cu, Ni) that serve as cofactors of many enzymes ^9^. Over 30% of the proteins found in living organisms bind metal ions, and a significant proportion of these proteins bind transition metals exclusively ^10,11^. The availability of metals may have shaped the evolutional trajectory of organisms ^12^. Some transition metals, such as Cd, are nonessential and toxic at relatively low concentrations, but can be taken up by plants via transporters for essential metals ^13^. Excess essential transition metals can also become toxic. Plants resist the toxic effects of excess transition metals mainly through chelation and compartmentation ^14^. An important class of metal-chelators are phytochelatins (PCs), oligopeptides synthesized from glutathione ^15^. Some transition metals can trigger immune responses in mammals ^16–22^. Both Zn^2+^ and Mn^2+^ bind to the DNA sensor cyclic GMP-AMP synthase (cGAS) and increase its production of the immune small-molecule messenger cyclic GMP-AMP (2′3′-cGAMP), enhancing the STING-dependent innate immune response ^21,22^. Zn^2+^ promotes condensate formation of cGAS-DNA, whilst Mn^2+^ directly promotes the activity of the cGAS enzyme. In addition, Mn^2+^ activates the NLR family pyrin domain containing 3 (NLRP3) inflammasome in mouse and human macrophages ^20^. Some transition metals can strengthen plant’s defense against pathogens with unknown mechanism ^23,24^; whether they can trigger plant immunity directly has not been reported.

Here, we show that a head-to-head gene pair of TNLs, STM1 and STM2, modulates transition metal-triggered immunity in an antagonistic manner. STM2 directly recognizes transition metals to activate immunity, which is suppressed by STM1. In *Arabidopsis*, tolerance to transition metal stress and resistance to bacterial wilt disease are balanced by the functional mode of this TNL pair.

## Results

### A pair of TNL genes are involved in the resistance to excess transition metals

The phytochelatin synthase-defective mutant *cad1-3* is Cd sensitive compared to wild type Col-0 due to the lack of PCs for internal detoxification ^25^. To explore the mechanism of Cd tolerance independent of the PCs pathway, we generated an EMS-mutagenized library of *cad1-3* and isolated a Cd hypersensitive mutant (*stm1-1 cad1-3*). Root and shoot growth of *stm1-1 cad1-3* was inhibited by increasing Cd concentration significantly more than *cad1-3* (Fig. 1a, b and Extended Data Fig. 1a-c). When grown in soil amended with moderate concentrations of Cd (1.0 and 2.5 mg kg^−1^), growth of *stm1-1 cad1-3* was also inhibited more than that of *cad1-3* (Fig. 1c and Extended Data Fig. 1d). The double mutant was also more sensitive to divalent transition metals including Zn, Cu(II), Co, Ni, Fe(II), Mn, and Pb, but not to other metals or metalloids including NaCl, arsenite, arsenate or Al (Fig. 1d), even though *cad1-3* was highly sensitive to arsenite and arsenate due to the role of PCs in arsenic detoxification ^26^. After we positionally cloned the causal gene, *STM1*, in the *cad1-3* background, we obtained T-DNA insertion lines of the gene (*stm1-2* and *stm1-3*) and generated a new knockout mutant by CRISPR/Cas9 (*stm1-4*) in the Col-0 background (Extended Data Fig. 1e,f). These single mutants were also more sensitive to divalent transition metals than Col-0, but not to alkaline-earth metals such as Ca and Mg (Fig. 1e). We named the mutants *sensitive to transition metals 1* (*stm1*). Higher doses of transition metals (except Fe^2+^) were used in the experiment with the single mutants because the phytochelatin synthase was functional. A dose of 70 µM Cd produced a similar degree of inhibition on root and shoot growth of the *stm1* single mutants to that of 30 µM Cd on the *stm1-1 cad1-3* double mutant (Extended Data Fig.1a-c and 1h-i). After exposure to 70 µM Cd for 1 day, the root and shoot cell saps of Col-0, *stm1-2*, and *stm1-4* contained 168-305 µM and 7.5-20.3 µM Cd, respectively, with no significant differences between genotypes (Extended Data Fig.1j-k). The majority of the Cd in the cell sap is likely to be complexed and stored in the vacuoles.

**Fig. 1:**
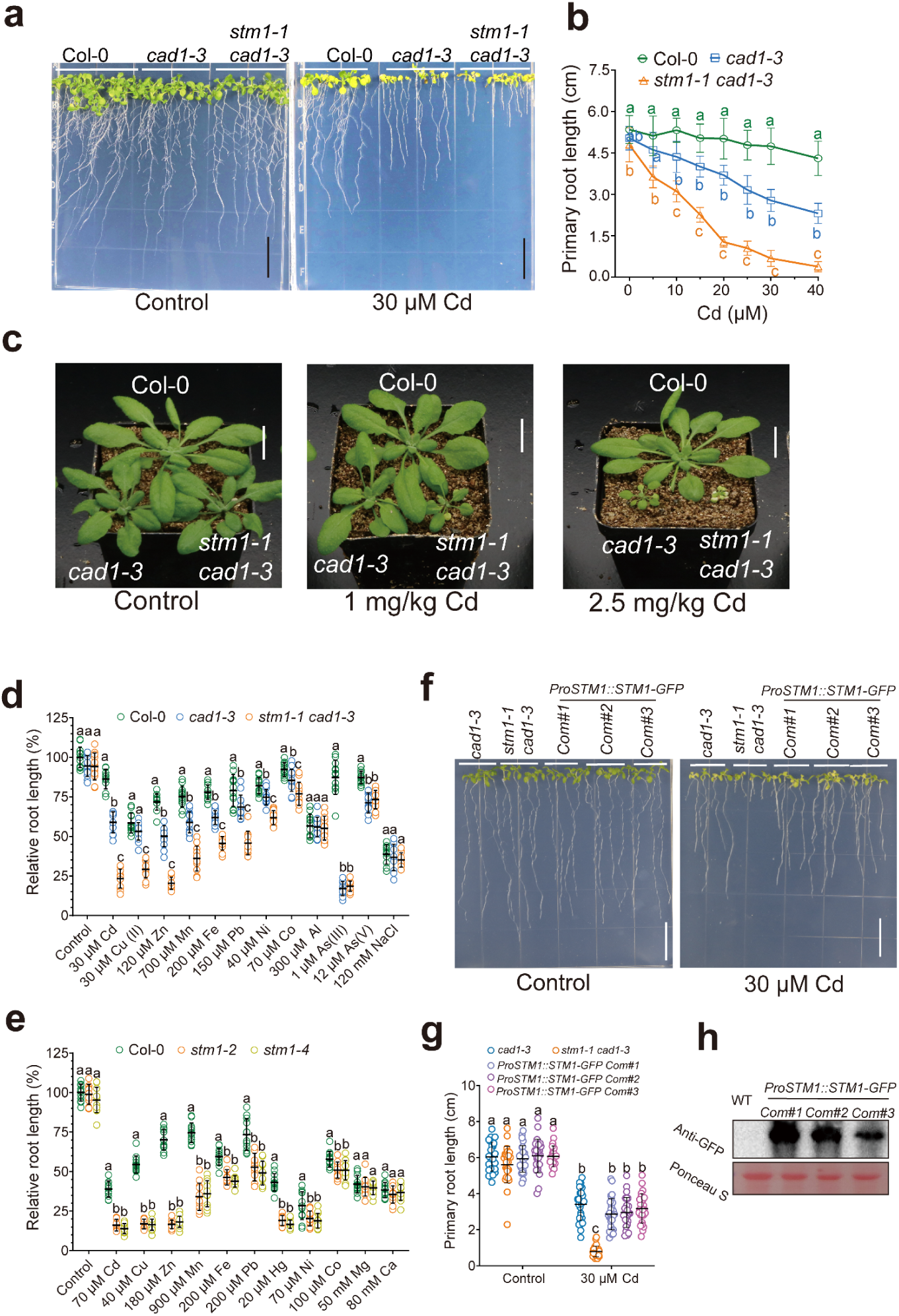
STM1 is involved in transition metal resistance in *Arabidopsis thaliana*. **a** Growth phenotypes of Col-0, *cad1-3*, and *stm1-1 cad1-3* grown in agar plates with or without Cd. **b** Response of primary root length of Col-0, *cad1-3*, *stm1-1 cad1-3* to increasing Cd concentration in agar plates. Data are means ± SD; n =12-15. **c** Growth phenotypes of Col-0, *cad1-3*, and *stm1-1 cad1-3* grown in soil amended with 0, 1.0, 2.5 mg/kg Cd. **d, e** Effects of various metals or metalloids on relative primary root length of Col-0, *cad1-3* and *stm1-1 cad1-3* (**d**) and of Col-0, *stm1-2* and *stm1-4* (**e**), expressed relative to that of Col-0 in the control. Lower concentrations of Cd, Cu, Zn, Mn, Pb, Ni, and Co were used for experiments in (**d**) than in (**e**) because of the loss-of-function of PCS in *cad1-3* and *stm1-1 cad1-3*. All data points are shown with means ± SD; n = 10-12.**f, g, h** Complementation of *ProSTM1::STM1-GFP* rescues the Cd sensitive phenotype of *stm1-1 cad1-3*; growth phenotype (**f**), primary root length (**g**), and detection of STM1-GFP in complementary lines by GFP antibody (**h**). All data points are shown in (**g**) with means ± SD; n = 24-28. Scale bars in (**a, c, f**) are 1.5 cm. Different letters represent significance at *P<*0.05 between genotypes within each time point (**b**), between each metal treatment (**d, e**) or among genotypes and Cd treatments (**g**) (one- or two-way ANOVA, followed by Tukey’s post-hoc multiple comparisons).

We generated a mapping population from the cross between *stm1-1 cad1-3* and *cad1-3.* Among the F_2_ progeny, 553 and 142 plants exhibited the Cd sensitive phenotypes of *cad1-3* and *stm1-1 cad1-3*, respectively, which is consistent with a 3:1 ratio (χ^2^ = 0.0054, *P* > 0.05), suggesting a single recessive mutation for the *stm1-1 cad1-3* phenotype. Using bulked segregant analysis combined with whole-genome resequencing, we mapped the causal gene to the 17.7 - 19.4 Mb region of chromosome 5. This region contains *PCS1* (*At5G44070*) and three candidate genes (*At5G44370*, *At5G45210*, *At5G46970*) with single nucleotide polymorphisms (Extended Data Fig.1l). Using complementation, we identified *At5g45210* as the causal gene for *stm1-1 cad1-3* (Fig.1f-h and Extended Data Fig.1m-n). The mutant allele has a single nucleotide change (G2617A) at the 3’ end of the third intron just before the fourth exon, leading to mis-splicing due to violation of the GU-AU rule and premature termination missing the last 96 amino acid residues (Extended Data Fig.1o). Knockout of *At5g45210* in either Col-0 or *cad1-3* background led to Cd sensitivity, further confirming *At5g45210* as the causal gene for *stm1 cad1-3* (Extended Data Fig.1p-s).

*STM1* is annotated as a member of the TNL disease-resistance gene family. The gene is located within a cluster of NLR genes that spans about 50 kb and includes the well-characterized pair, *RPS4/RRS1* (Extended Data Fig.2a). Next to *STM1* is another TNL gene (*At5g45200*) that is transcribed in the opposite direction from *STM1.* Given that many head-to-head located disease resistance genes work in pairs ^4–6^, we knocked out *At5g45200* in the *stm1-2* and in the *stm1-1 cad1-3* background (Extended Data Fig.2b). Knockout of *At5g45200* reverted the Cd sensitive phenotype of *stm1-2* to that of Col-0 (Fig 2a-b), and of *stm1-1 cad1-3* to that of *cad1-3* (Extended Data Fig.2c-d). In contrast, knockout of *At5g45200* in Col-0 did not affect Cd sensitivity (Extended Data Fig.2e-f). We hypothesized that this gene, which we named *STM2*, forms a pair with *STM1* to regulate tolerance to divalent transition metals. Phylogenetic analysis shows that *STM1* and *STM2* are clustered in two separate groups and fall among the nine head-to-head TNL pairs previously identified in *Arabidopsis* Col-0 ^27^ (Extended Data Fig.2g). STM1 is only about one half of the size of STM2 with a relatively short C-terminal LRR domain. STM1 and STM2 both lack an integrated domain (ID), which is often present and required for effector recognition by the sensor TNL in pairs ^28,29^. Knockout of *RPS4* or *RRS1* in *Arabidopsis* Col-0 did not affect Cd or Cu sensitivity in root growth assays (Extended Data Fig.2h-i).

**Fig. 2:**
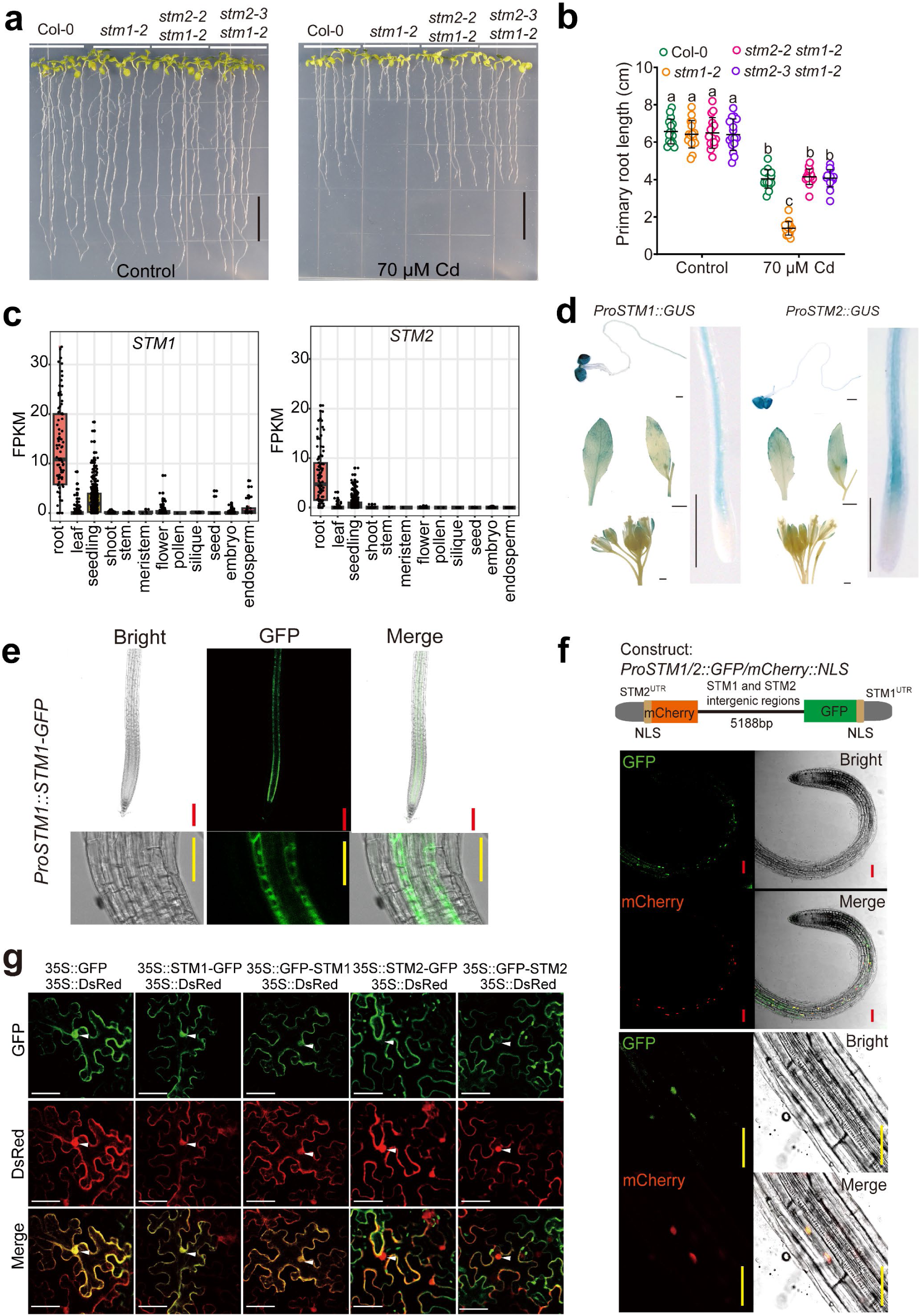
STM2 underlies the sensitivity to transition metals in *stm1*. **a, b** Cd sensitivity in *stm1-2* is rescued by knockout of *STM2* (see Extended Data Fig. 2b for *stm2* mutant alleles); growth phenotype (**a**) and primary root length (**b**). All data points are shown in (**b**) with means ± SD; n =16. Black bars in (**a**) = 1.5 cm. **c** *STM1* and *STM2* are mainly expressed in *Arabidopsis* roots. Transcriptomic data are from https://plantrnadb.com/athrdb/. FPKM, Fragments Per Kilobase of exon model per Million mapped fragments. **d** GUS staining of transgenic plants expressing *ProSTM1::GUS* or *ProSTM2::GUS*. Black bars = 500 µm. **e** Localization of STM1-GFP in the roots of transgenic plants expressing *ProSTM1::STM1-GFP.* Red bar = 100 µm, yellow bar = 50 µm. **f** Localization of STM1-GFP and STM2-mCherry in roots of transgenic plants expressing *ProSTM1/2::STM1-NLS-GFP/STM2-NLS-mCherry* plants (vector construction shown in the top panel). Red bar = 100 µm, yellow bar = 50 µm. **g** Subcellular localization of STM1 and STM2. *GFP*, *GFP-STM1*, *STM1-GFP*, *GFP-STM2*, and *STM2-GFP* were transiently expressed in *N. benthamiana* leaves driven by the CaMV 35S promoter. *35S::DsRed* was co-expressed as a marker for cytoplasm and nucleus (indicated by white arrowhead). White bar = 20 µm. Different letters in (**b**) represent significance at *P<*0.05 among genotypes and Cd treatments (two-way ANOVA followed by Tukey’s post-hoc multiple comparisons)

Analysis of more than one thousand *Arabidopsis* transcriptomes revealed that *STM1* and *STM2* are primarily expressed in the roots, with the former having a higher expression than the latter (Fig. 2c). We generated transgenic plants expressing *ProSTM1::GUS* and *ProSTM2::GUS* in Col-0 and *ProSTM1::STM1-GFP* in *stm1-1 cad1-3*. Histochemical staining of GUS showed that the two genes exhibited a similar expression pattern in the roots, leaves, and flowers (Fig. 2d). Exposure to Cd or inoculation of the bacterial wilt pathogen *Ralstonia solanacearum* to roots did not affect the expression of either *STM1* or *STM2* in the roots and shoots (Extended Data Fig.3a-h). Knockout of *STM2* did not affect *STM1* expression, nor did *STM1* knockout affect *STM2* expression (Extended Data Fig.3i-j). In the *ProSTM1::STM1-GFP* transgenic lines, the GFP was localized to a single layer of root cells, likely to be the endodermis (Fig. 2e). Moreover, STM2 and STM1 were colocalized in the endodermal cells, as revealed by transgenic plants expressing a construct comprising GFP and mCherry fused to a nuclear localization signal peptide at 5188 bp of their intergenic regions, followed by their respective 400 bp terminators (Fig.2f). Exposure to 70 µM Cd for 1 d did not change the spatial expression pattern of GFP and mCherry (Extended Data Fig.3k). In the root of *ProSTM1::STM1-GFP* transgenic lines, the GFP fluorescence was present in both the cytoplasm and the nucleus (Fig.2e). We could not detect GFP fluorescence in the transgenic lines of *STM2-GFP* driven by either CaMV35S or the native promoter of *STM2*. Instead, we transiently expressed *STM2-GFP* or *GFP-STM2*, and *STM1-GFP* or *GFP-STM1*, together with 35S:DsRed as a marker for nucleus and cytoplasm, in tobacco leaves. STM1-GFP or GFP-STM1 was present in both the nucleus and cytoplasm, while STM2-GFP or GFP-STM2 was present only in the cytoplasm (Fig.2g).

### Transition metals activate STM2-mediated immunity, which is suppressed by STM1

In the following experiments, we used Cd, as well as Cu and Zn in some experiments, as representative divalent transition metals. We performed RNA-seq analyses on the roots and shoots of *stm1-1 cad1-3* and *cad1-3* exposed to 0 or 30 µM Cd for 24 h. 1615 genes were upregulated by Cd in the roots of *stm1-1 cad1-3* compared with *cad1-3*. The differentially expressed genes are enriched in the immune function categories, such as “defense response to fungus”, “systemic acquired resistance”, and “response to chitin” (Extended Data Fig.4a-c). These categories converge on plant immune responses ^30,31^, and the Cd-induced transcriptional changes are similar to those observed in ETI^AvrRPS4 32^ (Extended Data Fig.4d). These results suggest that Cd triggers immune responses when *STM1* is not functional. In comparison, the transcriptome of the shoots was less affected (Extended Data Fig.4e-g).

RNA-seq analyses were also performed on Col-0, *stm1-2*, *stm2-2*, and *stm2-2 stm1-2* plants treated with or without 70 µM Cd for 16 h. Only *stm1-2* showed elevated expression of a range of immune function-related genes in response to Cd in the roots (Fig.3a and Supplemental Table S1). *PR2* and *PR5* are pathogenesis-related immune transcriptional markers ^33^; both genes were upregulated in the roots of *stm1-2* exposed to Cd, but not in *stm2-1* or *stm2-2 stm1-2* double mutant (Extended Data Fig.4h, i). Exposure of *cad1-3* to Cd up to 70 µM, which caused severe toxicity, did not upregulate *PR2* and *PR5* expression as did in *stm1 cad1-3* (Extended Data Fig.4j, k), indicating that upregulation of immune response genes by Cd in *stm1 cad1-3* is not an indirect effect of Cd toxicity. Consistent with the transcriptomic responses, we observed a significantly increased accumulation of both free and bound salicylic acid, a marker of plant immune metabolism, in the roots of *stm1-2* exposed to Cd (Fig.3b, c). Such responses were not observed in *stm2-1* or in the *stm2-2 stm1-2* double mutant.

**Fig. 3:**
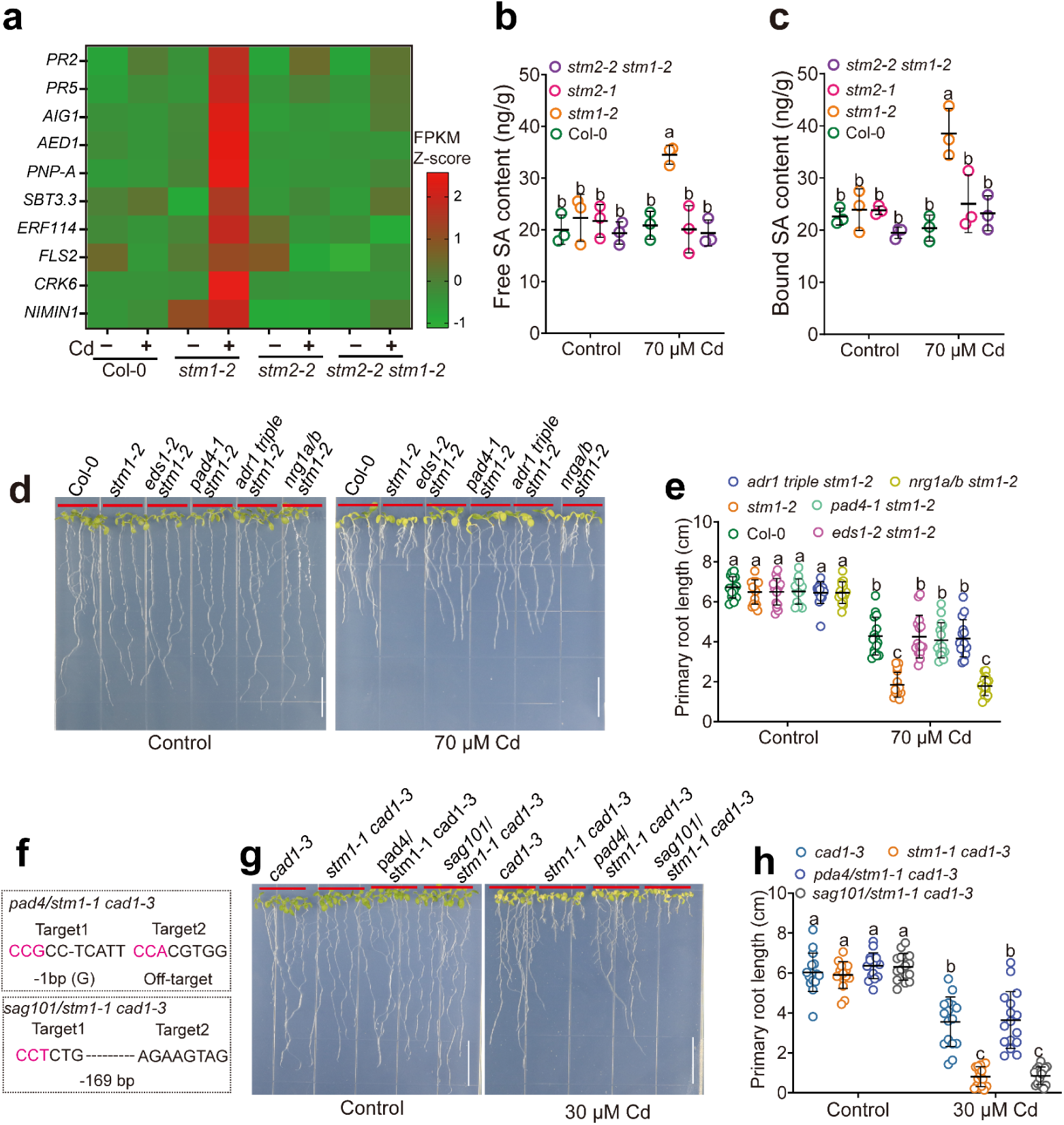
Cd induces STM2-activated immune responses in *stm1* via the EDS1-PAD4-ADR1 module. **a** Heatmap of ten representative immunity-related genes in roots of Col-0, *stm1-2*, *stm2-2*, *stm2-2 stm1-2* with or without 70 µM Cd for 16 h. Data are RNA-Seq Z-score normalized by the average expression levels of all genes in each sample. **b, c** Cd induced accumulation of free (**b**) and bound salicylic acid (SA) (**c**) in *stm1-2* roots, which was abolished by deletion of *STM2*. All data points are shown with means ± SD; n = 3. **d, e** Knockout of *EDS1*, *PAD4*, or *ADR1* rescued Cd sensitivity in *stm1*; growth phenotype (**d**) and primary root length (**e**). All data points are shown with means ± SD; n = 15. White bars = 1.5 cm. **f-h** Knockout of *PAD4*, but not *SAG101*, rescued Cd sensitivity in *stm1-1 cad1;3*; CRISPR/Cas9 edited mutant alleles of *PAD4* and *SAG101* in the *stm1-1 cad1-3* background (**f**); growth phenotype (**g**) and primary root length (**h**). All data points are shown with means ± SD; n = 16. Scale bars in (**d, g**) are 1.5 cm. Different letters in (**b, c, e, h**) represent significant difference at *P<*0.05 (two-way ANOVA, followed by Tukey’s post-hoc multiple comparisons).

In *Arabidopsis*, two immune signaling modules, EDS1-PAD4-ADR1 and EDS1-SAG101-NRG1, function downstream of TNL-mediated immunity to confer resistance and cause cell death ^34,35^. To examine which module is responsible for Cd-triggered immunity in *stm1*, we generated double or multiple mutants by crossing *stm1-2* with *eds1-2*, *pad4-1*, *adr1s* triple mutant, or *nrg1a/b* double mutant. Loss-of-function of either *EDS1*, *PAD4*, or *ADR1s* rescued the Cd sensitivity phenotype of *stm1-2*, whereas loss-of-function of *NRG1a/b* did not (Fig.3d, e). Furthermore, we used CRISPR/Cas9 to knock out *PAD4* and *SAG101* in the *stm1-1 cad1-3* background. Knockout of *PAD4*, but not *SAG101*, rescued the Cd sensitivity phenotype in *stm1-1 cad1-3* (Fig.3f-h). These results indicate that the Cd-induced STM2-dependent growth inhibition requires the EDS1-PAD4-ADR1s module.

To test whether STM1 or STM2 activates cell death, we performed hypersensitivity response (HR) assays using *Agrobacterium*-mediated transient expression in tobacco (*Nicotiana benthamiana*) leaves. Expression of *STM2*, but not *STM1*, triggered cell death, whereas co-expression of *STM1*, but not the truncated *STM1* cDNA from *stm1-1 cad1-3*, suppressed the STM2 activated HR (Fig. 4a and Extended Data Fig.5a-b,d-f). Addition of Cd (200 µM, similar to the root cell sap Cd concentration (Extended Data Fig.1j, k), 24 h after *STM2* expression significantly enhanced cell death, but did not cause cell death in the empty-vector (EV) control (Fig. 4b, Extended Data Fig.5c). The addition of Cd (200 µM) did not release the suppression by STM1 on STM2-activated HR (Extended Data Fig.5g-i). Reducing Cd concentration to 50 µM decreased the extent of cell death caused by STM2, but was still higher than the no Cd control (Extended Data Fig.5j-l). Similarly, application of Cu (100 µM) or Zn (400 µM) enhanced STM2-induced cell death (Extended Data Fig.5 m-o). The metals applied to tobacco leaves would be partially taken up by the leaf cells and undergo complexations and subcellular compartmentation, similar to that occurs in metal-exposed *Arabidopsis* plants.

**Fig. 4:**
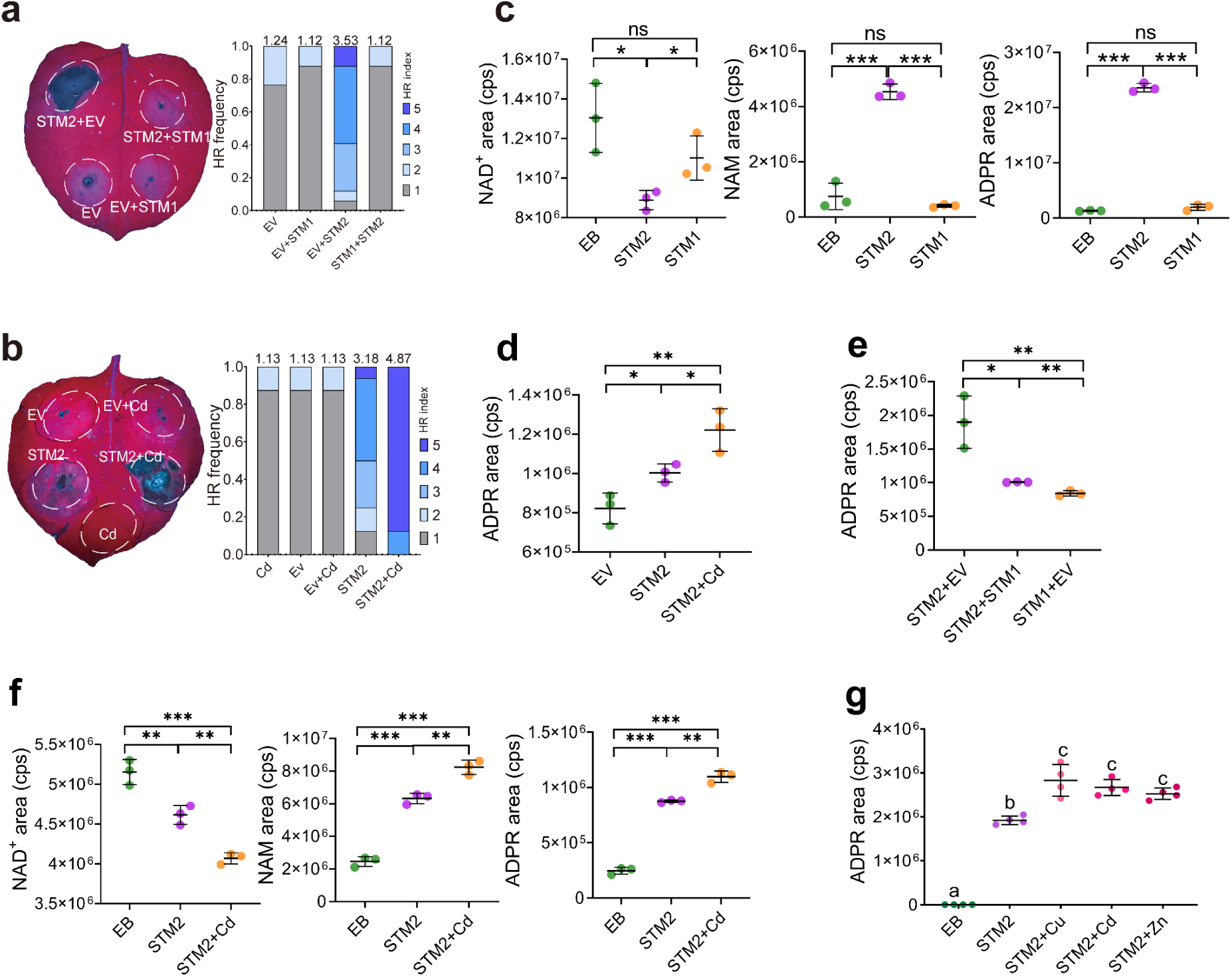
STM2-activated immune responses are suppressed by STM1 and promoted by transition metals. **a** Transient expression of *STM2* induced HR in tobacco leaves, which was suppressed by co-expression of *STM1*. Cell death phenotype under UV light (left) and frequency of HR score (right), n=17. EV, empty vector. HR index was scored according to Extended Data Fig.5a with weighted mean scores shown above the bars. Detections of STM2-YFP and Strep-STM1 fusion proteins in tobacco leaves are shown in Extended Data Fig.5b. **b** Cd enhanced HR caused by STM2. Cell death phenotype under UV light (left) and frequency of HR score (right), n=17. Cd (200 µM) or MES buffer was injected 24 h after infection with EV or *STM2*. Detections of STM2-YFP fusion protein in tobacco leaves are shown in Extended Data Fig.5c. **c** In vitro assay of the NAD^+^ hydrolytic activity of STM1 and STM2 recombinant proteins, measured by the depletion of the substrate NAD^+^ and the production of the products NAM and ADPR. EB, empty beads. All data points are shown with means ± SD; n = 3. **d** In vivo assay of the NAD^+^ hydrolytic activity of STM2 in tobacco leaves with or without Cd (200 µM), measured by the production of ADPR in leaf lysate. All data points are shown with means ± SD; n = 3. **e** In vivo assay of the NAD^+^ hydrolytic activity of STM2 in tobacco leaves with or without co-expression of STM1, measured by the production of ADPR in leaf lysate. All data points are shown with means ± SD; n = 3. **f** In vitro assay of the NAD^+^ hydrolytic activity of STM2 recombinant protein with or without Cd (50 µM), measured by the depletion of the substrate NAD^+^ and the production of NAM and ADPR. EB, empty beads. All data points are shown with means ± SD; n = 3. **g** In vitro assay of the NAD^+^ hydrolytic activity of STM2 recombinant protein with or without 50 µM of Cd, Zn, or Cu, measured by the production of ADPR. EB, empty beads. All data points are shown with means ± SD; n = 3. In (**c, d, e, f**),Significant difference at * *P*<0.05, ** *P*<0.01, and *** *P*<0.001 by two-sided *t*-test, in (**g**), different letters represent significant difference at *P<*0.05 (two-way ANOVA, followed by Tukey’s post-hoc multiple comparisons).

**Fig. 5:**
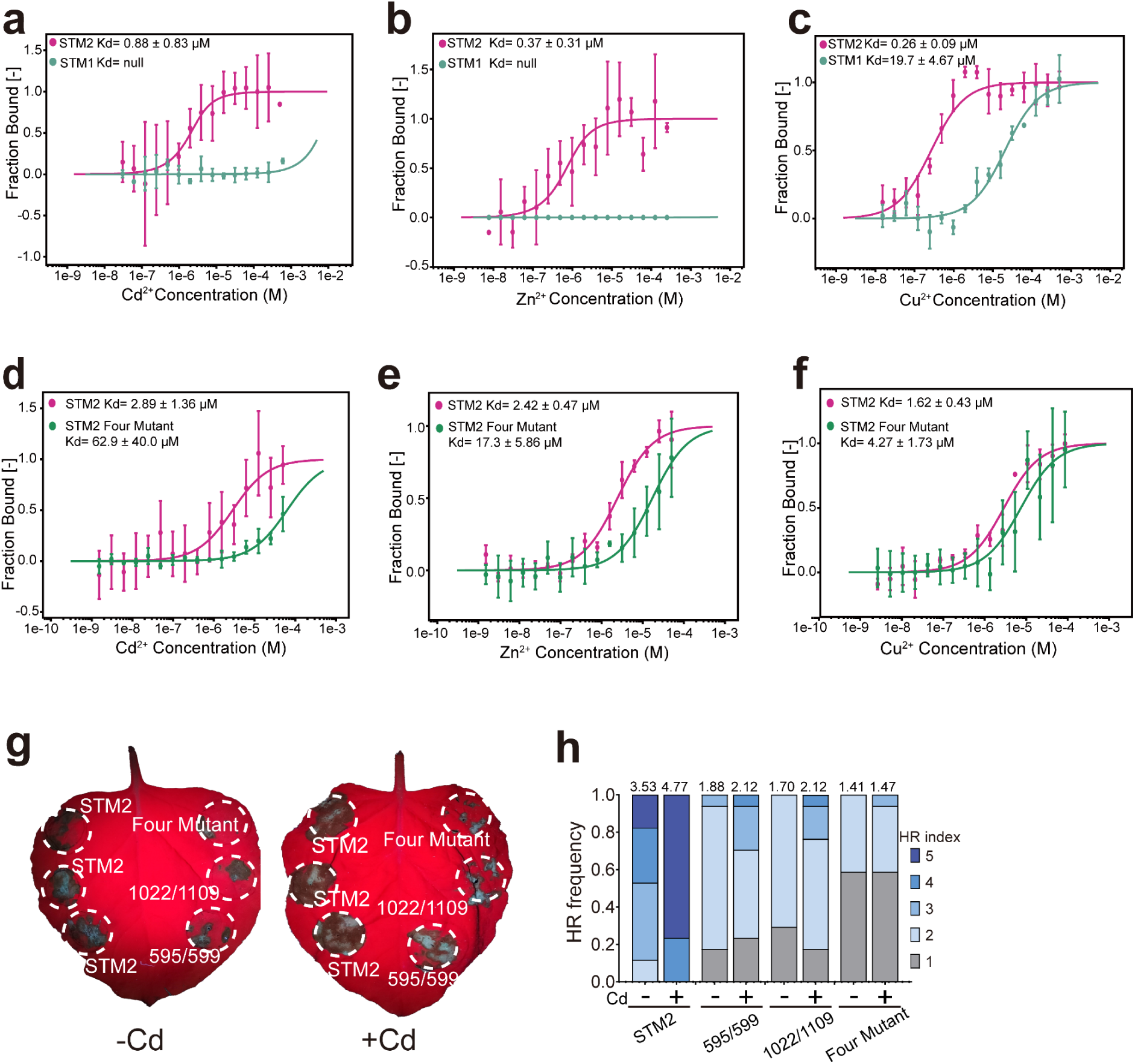
Binding of STM2 with Cd^2+^, Zn^2+^ and Cu^2+^. **a-c** MST assays of the binding of recombinant STM2 and STM1 proteins for Cd^2+^ (**a**), Zn^2+^ (**b**), and Cu^2+^ (**c**). **d-f** MST assays of the binding of recombinant wild-type STM2 and STM2-four mutant (C595, C599, H1022, and C1109 in the LRR domain are replaced with Ala) protein for Cd^2+^ (**d**), Zn^2+^ (**e**), and Cu^2+^ (**f**). **g, h** Tobacco HR assays of wild-type STM2 and mutated STM2 proteins in which C595/C599, H1022/C1109, or all the four amino acid residues were replaced with Ala without (lift) and with (right) Cd (200 µM); HR phenotype under UV light (**g**); Frequency of HR score with weighted means above the bars, n=17 leaves (**h**). Detections of STM2-YFP and mutated STM2-YFP fusion protein expressed in tobacco leaves by GFP antibody are shown in Extended Data Fig.8d.

To further investigate the immune signaling pathway downstream of STM2, we performed HR assays in the *N. benthamiana* quadruple mutant *epss*, which lacks the native *NbEDS1*, *NbPAD4*, *NbSAG101a*, and *NbSAG101b*. Expression of *STM2* did not induce HR in *epss* with or without Cd addition. In contrast, transient co-expression of *STM2* with *Arabidopsis AtEDS1*/*AtPAD4*/*AtADR1-L1* in *epss* leaves induced HR, which was markedly enhanced by the addition of Cd (Extended Data Fig.5p-r), whereas expression of *YFP* did not induce HR regardless of whether Cd was added (Extended Data Fig.5s-u). These results confirm that STM2 acts through the EDS1/PAD4/ADR1 module, and the effect of Cd requires the presence of STM2. In contrast to STM2, HR induced by *Arabidopsis* RPS4 and tobacco ROQ1 was not enhanced by Cd (Extended Data Fig.5v-x).

STM1 has a dual localization in the cytoplasm and the nucleus. We fused the nuclear localization signal peptide (NLS) or the nuclear export signal peptide (NES) to the C-terminus of STM1-GFP to modify its subcellular localization (Extended Data Fig.6a-b). STM1 suppressed cell death in tobacco leaves caused by STM2 when it was localized to the cytoplasm, but not in the nucleus (Extended Data Fig.6c). Furthermore, transgenic *Arabidopsis* plants expressing cytoplasm-localized STM1, but not nucleus-localized STM1, rescued the Cd sensitivity phenotype of *stm1-2* (Extended Data Fig.6d-f), implying that cytoplasmic localization of STM1 is required for its biological function of suppressing cytoplasm-localized STM2. Taken together, these results indicate that transition metals trigger immune responses through STM2, whereas STM1 acts as a suppressor of STM2.

### The NADase activity of STM2 is suppressed by STM1 but enhanced by transition metals

TNL proteins catalyze the cleavage of NAD^+^ to generate signaling molecules for immune activation ^36,37^. We expressed STM1 and STM2 in insect Sf9 or human HEK293T cells and purified the recombinant proteins. We found that STM2 cleaves NAD^+^ into nicotinamide (NAM) and ADP-ribose (ADPR) *in vitro*, whereas STM1 does not (Fig.4c). NAM and ADPR are typical NAD^+^ breakdown products and can be used to monitor the NADase activity of TNL protein^38,39^. The addition of Cd after the expression of *STM2* in tobacco leaves significantly increased the content of ADPR produced in the tobacco cell lysate (Fig.4d), whereas co-expression of *STM1* with *STM2* significantly reduced ADPR production (Fig.4e). We also determined the NAD^+^ cleavage activity of purified STM2 protein *in vitro* with or without Cd. The presence of Cd (50 µM) significantly promoted the hydrolysis of NAD^+^ and the production of NAM and ADPR (Fig.4f). Similarly, both Cu and Zn (50 µM) enhanced the hydrolysis of NAD^+^ to produce ADPR by STM2 *in vitro* (Fig.4g). These results suggest that transition metals enhance STM2 activation by promoting the NADase enzymatic activity of STM2, whereas STM1 suppresses the enzymatic activity of STM2.

The NADase activity of TNL proteins is catalyzed by the TIR domain with a conserved glutamic acid residue being essential for the enzyme activity^38,40^. A conserved glutamic acid (E91) is present in the TIR domain of STM2, but not in STM1 (Extended Data Fig.7a).

Expression of the TIR domain of STM2 in *E. coli* depleted cellular NAD^+^ as did the positive control RPS4-TIR, whereas expression of STM1-TIR did not (Extended Data Fig.7b). To examine the role of E91 in the Cd sensitivity of *stm1*, we edited the coding sequence of *STM2* in *stm1-2* to replace E at the 91^st^ position to glycine (G) or lysine (K) using a single-base editing technology (Extended Data Fig.7a) ^41^. Both substitutions restored the Cd sensitivity of *stm1-2* to the Col-0 level (Extended Data Fig.7c, d). In addition, STM2^E91G^, STM2^E91K^, and STM2^E91A^ lost the ability to cause cell death in tobacco leaves in HR assays (Extended Data Fig.7e-g) and the ability to produce ADPR (Extended Data Fig.7h). These results indicate that the NADase activity of STM2 is causally linked to its ability to cause HR in tobacco and the Cd sensitivity in *stm1*.

### STM2 binds transition metals

NLRs have been known to respond to pathogen effector proteins. The observation of Cd, Cu and Zn triggering STM2-dependent immunity and promoting STM2 enzymatic activity *in vitro* indicates that these divalent transition metals could function as a ligand to activate STM2 directly. Hence, we tested whether STM1 and STM2 can bind Cd^2+^, Zn^2+^ and Cu^2+^. Microscale thermophoresis (MST) assays showed that the recombinant STM2 could bind Cd^2+^, Zn^2+^ and Cu^2+^ with *K*_d_ values in the micromolar range (Fig.5a-c). In contrast, STM1 could not bind Cd^2+^ and Zn^2+^ but could bind Cu^2+^ with a much lower affinity (Fig.5a-c).

To explore the metal binding site in STM2, we expressed and purified the TIR and LRR domains separately and performed MST assays. The STM2 LRR domain binds Cd^2+^, Zn^2+^, and Cu^2+^ with *K*d values in the micromolar range, whereas the TIR domain binds the three metals with a much lower affinity (*K*d values 1-2 orders higher than those for the LRR domain) (Extended Data 8a). AlphaFold 3^42^ predicts that Zn and Cu ions are bound to C595, C599, C1109 and H1022 in the LRR domain of STM2 with a high confidence of prediction (Extended Data 8b, c); prediction of Cd binding is not available in AlphaFold 3. We used site-directed mutagenesis to replace the four amino acid residues with alanine in STM2. Compared with wild-type STM2 protein, the mutated protein showed a 22.0-, 7.1- and 2.6-fold increase in the *K*d value for Cd^2+^, Zn^2+^ and Cu^2+^, respectively (Fig.5d-f), indicating a marked decrease in the binding affinity. Tobacco HR assays showed that the ability to cause cell death was weakened markedly in the mutated STM2 proteins in which C595/C599, C1109/H1022, or all four amino acids were replaced with alanine. Moreover, Cd enhancement of HR was completely lost in the mutated STM2 in which the four amino acid residues were replaced with alanine (Fig.5g, h and Extended Data 8d). These data suggest that divalent transition metals are coordinated with C595, C599, C1109 and H1022 in the LRR domain and loss of these coordination sites diminishes the immune function of STM2.

### Homomeric and heteromeric interactions between STM2 and STM1

The oligomerization of NLR proteins is a prerequisite for their immune function^43–45^. Using bimolecular fluorescence complementation (BiFC) assay, we found that STM1 could physically interact with itself and with STM2 (Extended Data 9 a). Interaction between STM2 itself was also observed (Extended Data 9a). Transient co-immunoprecipitation (Co-IP) experiments for STM1 and STM2 conducted in tobacco leaves showed that, although STM2-YFP was expressed weakly, it was able to co-immunoprecipitate both Strep-STM1 and Strep-STM2, whereas the control YFP did not (Extended Data 9b). Strong immunoprecipitation between strep-STM1 and STM1-YFP was also evident (Extended Data 9b). Interactions between STM1 and STM2 were further verified by co-expression with the GFP and MBP tags in insect cells and GFP pull-down assay (Extended Data 9c). These results suggest that STM1 and STM2 can undergo both homomeric and heteromeric interactions, similar to the RPS4 and RRS1 pair ^44,45^.

We used gel filtration to examine whether STM1 affects multimerization of STM2. Flag-STM2 alone or Flag-STM2 + Strep-STM1 were separated by gel filtration. Eluate in different fractions were separated on SDS-PAGE. Flag-STM2 alone was eluted from 7 to 15 ml of eluate, corresponding to a minimum size of about 440 kDa, compared with approximately 148 kDa of the Flag-STM2 monomer (Extended Data 9d). This broad distribution pattern implies that STM2 protein may be in an unstable state of multimerization. The presence of Strep-STM1 with Flag-STM2 delayed the elution of the latter to 11 – 16 ml of eluate, corresponding to a minimum size of about 158 kDa, with co-elution of Strep-STM1 in the same eluate fractions, suggesting that STM1 hinders the multimerization of STM2, probably by forming STM1-STM2 complexes (Extended Data 9d). We propose that interaction between STM1 and STM2 hinders homomeric oligomerization of STM2 and thus inactivates its immune function. There was no evidence that Cd affects STM1-STM2 interaction in BiFC assays (Extended Data 10a) or STM2 oligomerization in BN-PAGE assays (Extended Data 10b).

In tobacco *epss* transiently expressing *STM2* with *AtEDS1/AtPAD4/AtADR1-L1*, BN-PAGE showed that Cd addition markedly enhanced the formation of high-order oligomers of ADR1-L1, likely also with EDS1 and PAD4 (Extended Data 10c). This is consistent with Cd-enhanced STM2 activity leading to enhanced ADR1-L1 oligomerization, triggering downstream immune responses.

### Trade-off between resistance to transition metals and bacterial wilt

*STM1* is *BWS1*, which was recently shown to confer bacterial wilt susceptibility via an unknown mechanism ^46^. To provide a mechanistic framework for these findings, we addressed the role of *STM1* and *STM2* in bacterial wilt resistance. We inoculated *Arabidopsis stm1* or *stm2* mutants and wild type (Col-0) with *Ralstonia solanacearum* strains GMI1000 or RS1115. Consistent with ^46^, Col-0 was susceptible to these bacterial wilt pathogen isolates, whereas *stm1* mutants were highly resistant (Fig.6a,b and Extended Data Fig.11a,b). Knockout of *STM2* in the *stm1-2* background increased the disease index to a level similar to that in Col-0 (Fig.6a,b and Extended Data Fig.11a,b), suggesting that the disease resistance in *stm1* is *STM2-*dependent. Knockout of *STM2* in the Col-0 background did not affect the disease index significantly, consistent with the immune function of STM2 being suppressed by STM1.

We further explored the interaction between Cd tolerance and *R. solanacearum* resistance. Plants were grown in Cd-free peaty soil for 4 weeks. The soil was then soaked with deionized water or 1 mM CdCl_2_ solution for 3 h and two days later inoculated with or without GMI1000. This relatively high concentration of Cd was used because the soaking period was brief, and the peaty soil had a high Cd adsorption capacity due to the high organic matter content (43%). The Cd concentration in the soil solution, the pool of Cd that is immediately available to plants, was 2.88 ± 0.72 µM, which is relatively low and environmentally relevant. In the absence of GMI1000 inoculation, *stm1-2* was more sensitive to Cd as the mutant grew significantly smaller shoot biomass than other genotypes and showed clear Cd toxicity symptoms (Fig. 6c-f). *stm1-2* was more resistant to GMI1000 inoculation than other genotypes in the absence of added Cd, but in the presence of the multiple transition metals typically present in the soil. This resistance was further significantly enhanced by the addition of Cd to the soil (Fig. 6c-f). The effect of Cd enhancing disease resistance was not observed in Col-0 or the *stm2-2 stm1-2* double mutants. After exposure to Cd for 4 days, roots of Col-0 and *stm1-2* contained 151-334 µM Cd in the cell sap (Extended Data Fig.11c); these concentrations were similar to those in the agar plate experiment exposed to 70 µM Cd for 1 day (Extended Data Fig.1j), suggesting comparable levels of Cd exposure. Experiments with Cu showed similar responses (Extended Data Fig.11d-g). These results indicate that transition metals enhance, whereas STM1 suppresses, the immunity activated by STM2 and that there is a trade-off between sensitivity to transition metals and susceptibility to *R. solanacearum* (Fig.6g).

**Fig. 6:**
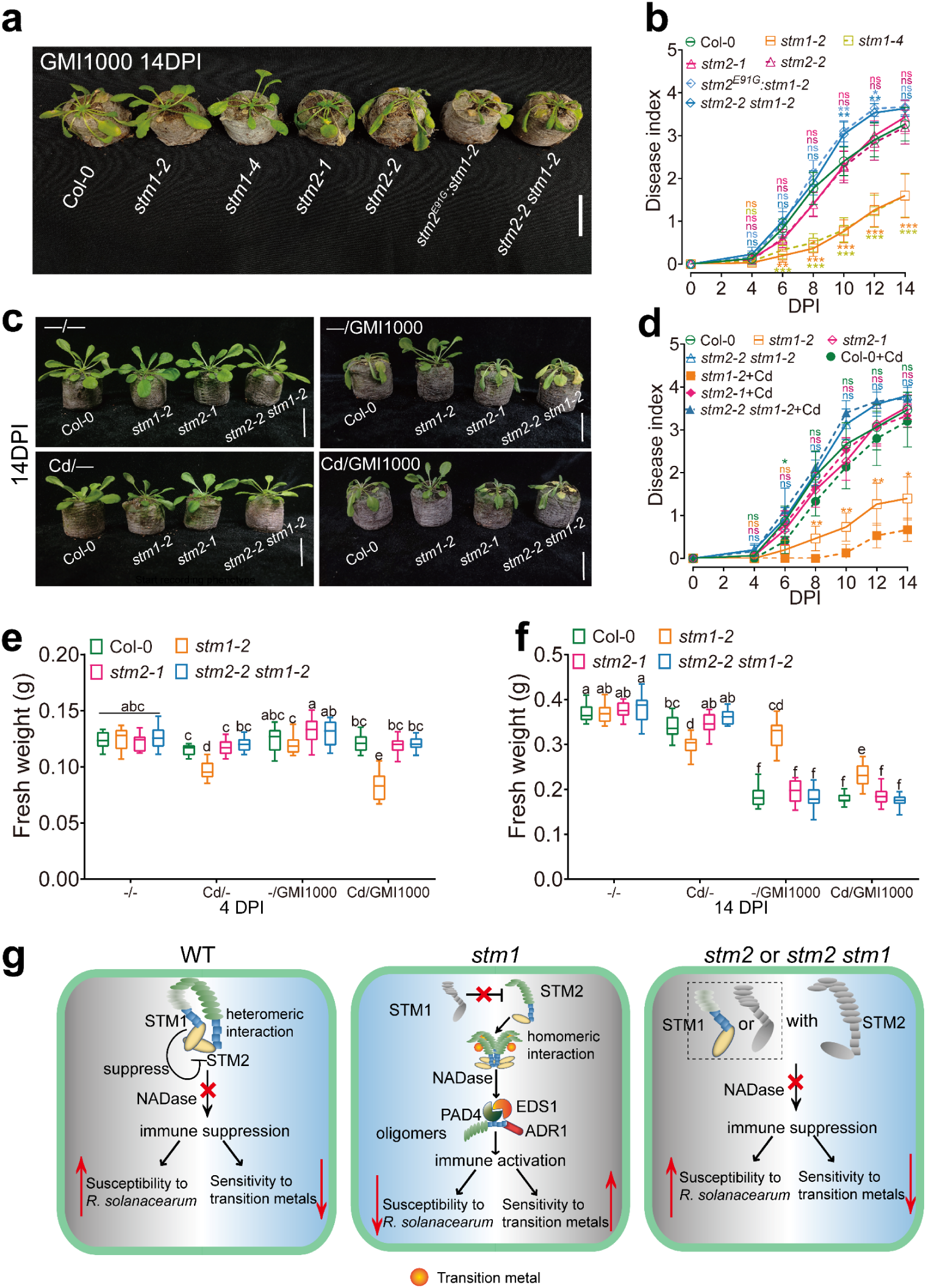
STM1 and STM2 mediate the trade-off between transition metal sensitivity and susceptibility to bacterial wilt disease. **a, b** Mutations in *STM1* enhanced resistance of *Arabidopsis thaliana* to *R. solanacearum* strain GMI1000, but the resistance was lost by the deletion of *STM2*; bacterial wilt phenotype (**a**) and disease index after pathogen infection (**b**). *stm2^E91G^:stm1-2* is a single base edited line of *STM2* in the *stm1-2* background. Scale bar = 3.5 cm. **c-f** Interactions between *STM1/STM2* genotype and Cd on resistance to *R. solanacearum* strain GMI1000; plant growth and bacterial wilt phenotype (**c**), disease index (**d**), plant biomass at 4 DPI (**e**) or 14 DPI (**f**). Scale bar = 3.5 cm. **g** Proposed model of STM1-STM2 in trade-off between sensitivity to transition metals and susceptibility to bacterial wilt pathogen. Data in (**b, d**) represent means with 95% confidence interval, n = 15 biological replicates in (**b**); n= 30 biological replicates for 0 – 4 DPI and 15 biological replicates for 6 – 14 DPI in (**d**). In the boxplots (**e**) and (**f**), the central line represents the median, the box 25-75 percentiles and the whiskers 5-95 percentiles.Statistical analysis was performed to compare the difference from Col-0 in (**b**), and between +/- Cd in the same genotype in (**d**) by two-sided *t*-test; ns, not significant, *\* P*<0.05, ** *P*<0.01, *** *P*<0.001, and among all. genotype/Cd/inoculation combinations in (**e, f**) by three-way ANOVA followed by Tukey’s post hoc tests. Different letters in (**e, f**) indicate significant difference at *P*<0.05.

## Discussion

In this study, we uncover a new functional model of NLRs in which the head-to-head paired NLR STM1 and STM2 play opposite roles in the resistance of *Arabidopsis* to transition metals and the soil-borne disease *R. solanacearum*, a bacterial wilt pathogen that causes havoc to many crops (Fig.6g). The co-localization of STM1/STM2 specifically in the root endodermis is congruent with the endodermis being a barrier for both metal ions and soil-borne pathogens ^47,48^. It has been shown that the endodermis of *Arabidopsis* roots accumulate high concentrations of transition metals including Fe, Zn and Mn ^49^. Unlike STM1/STM2, most *Arabidopsis* NLRs are mainly expressed in the leaves and, until now, no functional NLRs localized specifically in root cells have been characterized^50^. We found that STM2 is the executor of immunity via its NAD^+^ hydrolytic activity and is required for activation of the downstream EDS1-PAD4-ADR1s signaling module. STM2 directly binds Cd^2+^, Zn^2+^ and Cu^2+^ and likely also other transition metals, which in turn enhances its enzymatic activity. We found that three cysteine and one histidine residues in the LRR domain of STM2 are involved in the binding of transition metal ions; cysteine and histidine are well known for their affinities for transition metals^12^. Mutagenesis of these residues not only weakens the binding of transitions metals markedly but also abolishes Cd-enhanced HR by STM2. The *K*d values in the micromolar range, measured in in vitro assays, suggest that STM2 has relatively low affinities for the transition metal ions; such *K*d values are not uncommon for metalloproteins requiring metal ions for regulatory functions^51^. Metal carriers or chaperones may be required to deliver the metal ions to the target proteins^51^. Given that transition metals do not appear to interact directly with the catalytic TIR domain and do not directly promote oligomerization of STM2, how does binding of transition metals with the LRR domain enhance the NADase activity of STM2? It has been shown that the NB-LRR domains of flax TNL L6 negatively regulates the TIR domain via intramolecular interactions in the resting state^52^. We propose that the binding of the STM2 LRR domain with transition metals may release the negative regulation of LRR domain over the TIR domain, thus enhancing its NADase activity. Future studies are required to reveal how transition metals enhance the NADase activity of STM2.

In contrast, STM1 is a suppressor of STM2 likely through its heteromeric interactions with STM2, preventing oligomerization of STM2. STM1 protects plants from the growth inhibition caused by transition metal-triggered immunity and consequently dampens the resistance to bacterial wilt disease (Fig.6g). Exposure to Cd did not relieve the inhibition of STM1 on STM2, likely because the interaction between STM1 and STM2 is strong. Although the *Ralstonia* effector RipAC can also interact with BWS1/STM1 in vitro ^46^, the fact that the presence of STM1 suppresses the resistance to *Ralstonia* suggests that the interaction of RipAC and STM1 also does not relieve the inhibition of STM1 on STM2 and thus does not activate the immune function of STM2. In the absence of STM1, transition metals boost disease resistance by enhancing the activity of STM2. The STM1-STM2 model reveals a trade-off between metal sensitivity and disease susceptibility. STM1 may have evolved to suppress STM2-induced metal sensitivity at the expense of disease resistance because transitions metals are ubiquitous and pervasive in soil. The necessity to overcome metal sensitivity may therefore constitute a limiting factor for plant resistance against soil-borne pathogens.

Natural variation in *BWS1/STM1* among different *Arabidopsis* accessions was found to correlate with resistance to *R. solanacearum*^46^, suggesting that there exists natural variation in the function of BWS1/STM1. We propose that weak alleles of *STM1* would be less able to antagonize STM2 and hence benefit disease resistance. However, strong alleles of *STM1*, such as that in the Col-0 accession used in the present study, may be required to suppress unwanted immunity triggered by transition metals in soils with a high availability of these metals. It would be interesting to evaluate the relationship between different haplotypes of STM1 and sensitivity to transition metals versus resistance of bacterial wilt. Similarly, the presence or absence of *ACQOS/VICTR* among natural accessions of *Arabidopsis* provides a trade-off between resistance to disease resistance or to osmotic stress. The *ACQOS/VICTR* gene was found to be absent in 72% of the accessions tested, suggesting that resistance to osmotic stress takes priority over disease resistance ^8^.

In contrast to STM2, Cd did not enhance the HR caused by RPS4 or ROQ1. Mutations in *RPS4* or *RRS1* also did not lead to Cd sensitivity in *Arabidopsis*. Unlike *STM1* and *STM2*, the *RPS4-RRS1* pair is mainly expressed in leaves, where the concentrations of divalent metals are generally much lower than those in root. Also, unlike STM2, RPS4 and ROQ1 do not possess four strong metal binding residues (Cys and His) in their LRR domain according to the predictions by AlphaFold 3, which may explain their lack of HR response to Cd. Whether there are other transition metal-binding NLRs remains to be investigated.

## Methods

### Plant materials and growth conditions

*Arabidopsis thaliana* materials are listed in the Supplemental Table S2. For metal toxicity assays, gene expression and determination of SA, plants were grown on agar (1% w/v, Sigma-Aldrich, USA)-solidified plates with ½ Murashige-Skoog (MS) basal salt mixture (Hopebiol, HB8469) and 1% sucrose. Agar plate assays were conducted in a controlled environment chamber with 16 h daylength, 80–100 μmol m^−2^ s^−1^ light intensity, 22 °C and 65% relative humidity. *R. solanacearum* infection assays were conducted in a controlled environment chamber with 12 h daylength, 80–100 μmol m^−2^ s^−1^ light intensity, 28 °C and 75% relative humidity. *Nicotiana benthamiana* was grown in a greenhouse with 12 h daylength, 80–100 μmol m^−2^ s^−1^ light intensity, 26°C/22°C day/night temperature, and 60% relative humidity.

### Mutant screening and identification

An ethyl methanesulfonate-mutagenized library in the *cad1-3* mutant background was generated as described previously^54^. M_2_ seeds were germinated and grown for 12 d vertically on ½ MS agar plates containing 30 μM Cd, which inhibited root growth of *cad1-3* by approximately 50%. Plants showing more severe Cd-induced inhibition of root growth than *cad1-3* were rescued and grown to maturity for the collection of M_3_ seeds. The Cd-sensitive phenotype was verified using M_3_ seeds. A mutant (*stm1-1 cad1-3*) was selected for further characterization.

### Metal toxicity assays

*Arabidopsis* seeds were surface sterilized with 8% sodium hypochlorite for 10 min and stratified at 4 °C for two d. Seeds were geminated on ½ MS agar plates with or without additions of various metals or metalloids, including Cd (CdCl_2_), Cu (CuCl_2_), Zn (ZnCl_2_), Co (CoCl_2_), Ni (NiCl_2_), Fe (FeSO_4_), Hg (HgCl_2_), Pb (PbSO_4_), Al (AlCl_3_), NaCl, Ca (CaCl_2_), Mg (MgCl_2_), Na_3_AsSO_4_, or NaAsSO_4._The concentrations of the metals or metalloids used were based on results from preliminary range finding experiments. Plants were grown for 12 d before roots were photographed and root length quantified using Fiji-ImageJ2 (https://fiji.sc/). In the Al toxicity assay, pH of the growth medium was lowered to 4.5. Plants were also grown to maturity in the soil (peat-vermiculite mixture, pH 5.6, organic matter content 7.2%). Cd (as CdCl_2_) was added to the soil at 0, 1 or 2.5 mg Cd kg^−1^, and plants were grown under 12 h daylength for 8 weeks. Plants were photographed at different time points and rosette diameters were measured using Fiji-ImageJ2. The concentrations of 0.01 M CaCl_2_-extractable metals are shown in Supplemental Table S3.

### Map-based cloning of *STM1*

To identify the causal gene for the Cd sensitivity phenotype of *stm1-1 cad1-3*, we generated an F_2_ mapping population by crossing *cad1-3* with *stm1-1 cad1-3*. The F_2_ plants that showed a root growth inhibition phenotype by 30 µM Cd similar to that in *stm1-1 cad1-3* were selected for DNA extraction for whole-genome resequencing using the MutMap method ^55^. Briefly, DNA was extracted from 70 F_2_ individuals and mixed in an equal ratio. DNA of *cad1-3* plants was used as the control. A DNA library (10 μg mixed DNA) was prepared for Illumina sequencing according to the protocol for the Paired-End DNA Sample Prep kit (Illumina). Whole genomic resequencing was performed with a mean coverage of 40× using a paired-end read length of 150 bp on an Illumina HiSeq4000 sequencer. Sequence reads were mapped to the *Arabidopsis* Col-0 reference genome (TAIR 10) using BWA software ^56^. Three candidate genes were identified by MutMap. Derived cleaved amplified polymorphic sequence markers were developed to determine the co-segregation of the Cd sensitivity phenotype with SNPs (single nucleotide polymorphisms) in these candidate genes. The causal gene was identified based on transgenic complementation.

### Generation of complementation transgenic lines

Genetic complementation lines of *At5G44370*, *At5g45210*, *At5g46970* were generated in the *stm1-1 cad1-3* background. The DNA fragments of the three genes were amplified by PCR from the Col-0 genomic template containing promoters of 1908 bp, 2718 bp, and 758 bp, genomic sequences of 1299 bp, 2914 bp, and 495 bp, and 3’UTR sequences of 896 bp, 902 bp, and 364 bp after the stop codon, respectively. The DNA fragments were cloned into the plant expression vector pZH2B using the ClonExpressII One Step Cloning Kit (Vazyme, C112-01). A 2718 bp *STM1* promoter, 2091 bp *STM1* CDS sequence (Locus ID: *At5g45210.1*, no stop codons), eGFP and the NOS terminator sequence fragments were cloned into pSAT6A-EGFP-N1 and ligated sequentially into the pZH2B vector by using the ClonExpress MultiS One Step Cloning Kit (Vazyme, C113-01). Nucleic acid fragments of NES or NLS signaling peptides were synthesized and ligated to the C-terminus of eGFP by PCR. The fragment was combined with the C-terminal of *STM1* CDS through multi-fragment recombination and joined together using the pMDC32 vector (35S promoter and NOS terminator) which were linearized by *Kpn* I and *Hind*III. These constructions were transformed into *stm1-1 cad1-3* using the *Agrobacterium tumefaciens* (strain GV3101)-mediated floral dip method ^57^. More than 10 transgenic lines per construct were screened on ½ MS agar plates containing 50 μg/ml of hygromycin. T_3_ homozygous transgenic plants were used in phenotype assays.

### Tissue and subcellular localization of STM1 and STM2

Transcriptomic data sources (https://plantrnadb.com/athrdb/) ^58^ were mined to determine differential tissue expression of *STM1* (Locus ID: *At5g45210.1*) and *STM2* (Locus ID: *At5g45200.1*). To examine the subcellular localization of STM1 and STM2, the CDS of *STM1* (*At5g45210.1,* 2091 or 2094 bp, without or with the stop codon) and *STM2* (*At5g45200.1,* 3783 or 3786 bp, without or with the stop codon) were cloned into pSAT6A-EGFP-N1 and pSAT6-EGFP-C1 initial vectors which were linearized by *BamH* I and *Hind*III. These fusion constructs were introduced into pRCS2-ocs-nptII final vector. All expression constructs are listed in the Supplemental Table S4. These constructs, as well as pRCS2-ocs-nptII containing 35S::GFP, were co-expressed with *pRCS2-ocs-nptII* containing 35S::DsRed (a fluorescence marker for cytoplasmic and nuclear localization) in tobacco leaves using an *Agrobacterium*-mediated transient expression method ^59^. Images were taken using a confocal microscope (SP8, Leica). GFP fluorescence was observed at 496 nm for emission and 488 nm for excitation, and DsRed fluorescence at 584 nm for emission and 555 nm for excitation.

To examine the tissue expression pattern, a 2718 bp *STM1* promoter and a 2913 bp *STM2* promoter were ligated into the pCAMBIA-1300-GUS (NOS terminator) vector. The constructs were transformed into Col-0 by the *Agrobacterium*-mediated floral dip method ^57^. More than 8 transgenic lines each were screened on ½ MS agar plates containing 50 μg/ml of hygromycin. Eight-day-old plants of *ProSTM1::GUS* and *ProSTM2::GUS* transgenic lines and flowers at the anthesis stage were used for GUS staining. Following overnight incubation in a GUS staining solution, samples were washed twice with distilled water and photographed under a stereomicroscope (OLYMPUS MVX10).

The head-to-head pair of *STM1* and *STM2* shares an intergenic region of 5,188 bp. To examine whether STM1 and STM2 are co-localized in the same tissues, the intergenic region was cloned from the Col-0 genomic template via PCR and inserted into the vector pZH2B between the *Xba* I and *Kpn* I sites. The pZH2B:5188 bp vector was linearized using *Kpn* I restriction enzyme and ligated to the eGFP-NLS and 491 bp STM1 3’UTR tandem fragments by using the ClonExpress MultiS One Step Cloning Kit (Vazyme, C113-01). The generated STM1^UTR^-eGFP-NLS-pZH2B:5188 bp vector was linearized by *Xba* I, and the linear fragment was ligated to the mCherry-NLS and 776 bp STM2 3’UTR tandem fragments to generate the construct as shown in the schematic diagram in Fig. 2f. The construct was transformed into the Col-0 background. Eight-d-old transgenic plants were stained in Calcofluor White M2R (4193-55-9, Sigma) at a final concentration of 1 g l^−1^ for 30 min, washed twice with 0.01M PBS (pH=7.0), and the roots were observed under a confocal laser scanning microscope (SP8, Leica) at 335 nm, 496 nm and 610 nm for emission, respectively, and 433nm, 488 nm and 587 nm for excitation, respectively, for the fluorescence of calcofluor white, GFP and mCherry.

### Generation of gene-edited lines and high-order mutants

We used CRISPR/Cas9 to knockout *STM1* in the Col-0 and *cad1-3* backgrounds, to knockout *STM2* in the Col-0, *stm1-2* and *stm1-1 cad1-3* backgrounds, and to knockout *PAD4* or *SAG101* in the *stm1-1 cad1-3* background. Knockout targets were designed by using CRISPR-GE (http://crispr.hzau.edu.cn/)^60^. pYLCRISPR/Cas9-DH^61^ was employed to knock out *STM1* in the Col-0 and *cad1-3* background. pYAO:CRISPR/Cas9^62^ was employed to knock out *STM2* in the Col-0, *stm1-2* and *stm1-1 cad1-3* background, and to knock out of *PAD4* or *SAG101* in the *stm1-1 cad1-3* background. More than 60 transgenic plants were obtained for each line. T_2_ homozygous lines were identified using PCR and Sanger sequencing.

Single base editing technology was used to modify the codon for the glutamate residue at the 91^st^ position in the *STM2* coding sequence in the *stm1-2* background. The CBEmax-nCas9NG-p19 and ABEmax-nCas9NG-p19 system used has relaxed PAM preferences, resulting in the conversion of C·G and A·T pairing to T·A and G·C respectively ^41^. ‘GATTGCAGCTCATTTAAGCACCAT’ and ‘AAACATGGTGCTTAAATGAGCTG’ nucleotides were synthesized and formed in vitro at 95℃ for 5 min and 4℃ for 5 min into complementary paired double-stranded complexes, which were introduced by T4 ligase into CBEmax-nCas9NG-p19 vector linearized by *Bsa* I. The ‘GATTGATGAGCTGGTAAAGATCAA’ and ‘AAACTTGATCTTTACCAGCTCATC’ complexes were introduced into ABEmax-nCas9NG-p19 vector using the same method. The resulting constructs were transformed into *stm1-2* plants. More than 100 transgenic plants were obtained per line. The presence of desired mutations in T_1_ plants was determined by PCR. Homozygote lines were identified in the T_2_ generation.

To generate high-order mutants, *stm1-2* was crossed with *pad4-1*, *eds1-2*, *nrg1a/b*, or *adr1 triple*^63–66^. Gene accession numbers are shown in Supplemental Table S5. The homozygous high-order mutants were verified by PCR or by Sanger sequencing. More than 5 independent inbred lines were obtained for each high-order mutant.

### Phylogenetic analysis

We used the tBLASTp algorithm in the NCBI database to obtain homologous sequences of STM1 and STM2, retaining only the *Arabidopsis* TNL-like proteins. The full-length sequences of these proteins were aligned with ClustalW and a phylogenetic tree was constructed using the maximum likelihood method in MEGAX.

### RNA extraction and transcriptomic analysis

*cad1-3* and *stm1-1 cad1-3* plants were grown on 1/2 MS agar plates for 12 d and then transferred to plates with or without 30 μM Cd for 24 h. Col-0, *stm1-2*, *stm2-2*, *stm2-2 stm1-2* plants were grown on 1/2 MS agar plates for 12 days and then transferred to plates with or without 70 μM Cd for 16 h. More than 150 plants from 5 plates of each line in the same treatment were collected as one biological replicate, with 3 replicates per line and treatment. Roots and shoots were collected separately. Total RNA was extracted using a plant Total RNA Extraction Kit (BioTeke) following the manufacturer’s protocol. One μg RNA per sample was used for construction of cDNA library using the NEBNext® Ultra™ II RNA Library Prep Kit for Illumina® according to the manufacturer’s instructions. The products were purified (AMPure XP system) and quantified using the Agilent high sensitivity DNA assay on a Bioanalyzer 2100 system. The cDNA libraries were sequenced on NovaSeq 6000 platform (Illumina), generating >39 M raw 150 bp paired end reads per sample. The raw data were cleaned using Fastp (0.22.0) software. Filtered reads were compared to the reference genome (TAIR10.1, https://www.arabidopsis.org/) using HISAT2 software. Fragments Per Kilobase of exon model per Million mapped fragments (FPKM) and the read counts of each gene were calculated and obtained by RESM software. Differentially expressed genes (DEGs), defined as those with |log_2_FoldChange| > 1 and significance *P*-value < 0.05, were analyzed using DESeq2. Bi-directional clustering was performed to obtain gene expression heatmaps using the R pheatmap package. The heatmap was horizontally divided into nine gene sets as clusters, and each cluster was subjected to trend analysis and GO enrichment analysis. Based on the hypergeometric distribution, the top six GO entries with the lowest *P*-values were presented, with Cd-induced changes in the gene Cluster being highlighted. Col-0, *stm1-2*, *stm2-1*, *stm2-2 stm1-2* plants were grown on 1/2 MS agar plates for 12 d and then transferred to plates with or without different concentrations of Cd for 24 h. In the experiment with *R. solanacearum* infection, Col-0 plants were grown in Jiffy pellets for 4 weeks and then inoculated with bacterial strain GMI1000 for 1 d. Roots and shoots were collected for transcript analysis. Expression of *STM1*, *STM2, PR2,* and *PR5* were determined by quantitative RT-PCR performed on a CFX96 Touch Real-Time PCR Detection System (Bio-Rad) using the ChamQ SYBR qPCR Master Mix Kit (Q311-02, Vazyme). *UBQ10* and *Actin2* were used as the reference genes. All primers used in this study are listed in Supplemental Table S6. The RNA-seq data are deposited in NCBI GEO database (GSE291114 for *cad1-3* and *stm1-1 cad1-3*; GSE291226 for Col-0, *stm1-2, stm2-2*, *stm2-2 stm1-2*).

### Determination of free and bound salicylic acid (SA)

Col-0, *stm1-2*, *stm2-1*, and *stm2-2 stm1-2* plants were grown on 1/2 MS agar plates for 10 days and then transferred to plates with or without 70 μM Cd for 24 h. Roots and shoots were collected separately and stored at -80 °C prior to analysis. Extractions of free and bound SA (i.e. inactive SA in the form SA-2-O-β-D-glucoside and SA-β-D-Glucose Esterglucose ester stored in vesicles) were performed as described previously ^67^. In brief, 0.05 g of tissue samples were powdered in liquid nitrogen and extracted first with 70% ethanol and then with 90% methanol. Both extracts were combined and 0.1 volume of 20% trichloroacetic acid was added, followed by the addition of 1 volume of ethyl acetate/cyclohexane (1:1 v/v). The content was vortexed for 30 s and centrifuged at 10,000 *g* for 10 min at room temperature. The upper organic phase, containing free SA, was transferred to a 2 ml microcentrifuge tube and the procedure was repeated once and combined. The lower aqueous phase was hydrolyzed by adding 4 M HCl at 80 °C to release the bound SA and then extracted in the same way. Both extracts were evaporated by nitrogen gas blowing and dissolved in 100 µl methanol for HPLC analysis. The concentrations of SA were determined by HPLC with a C18 HPLC column. The mobile phase was a mixture of 0.2 M sodium acetate and MeOH (9:1, v/v, pH=5.5). The flow rate was 1 ml min^−1^ and the column temperature was maintained at 30 °C. The SA peak appeared at 9.5 min. The fluorescence of SA was detected at 305 nm. The concentration of SA was quantified by external calibration.

### Site-directed mutagenesis

*STM2^E91A^*, *STM2^E91K^*, *STM2^E91G^*, mutations of *STM2* with C595A, C599A, C1109A and H1022A, and mutations in the p-loop motif of *STM2* (G247A, G249A, K250A, T252A) were introduced using the Q5 site-directed mutagenesis kit (E0554S, NEB). All constructs were verified by DNA sequencing.

### Hypersensitive response (HR) assays

All expression constructs used in HR assays were constructed using the Gateway cloning system and listed in Supplemental Table S4. *Agrobacterium tumefaciens* strain GV3101 harboring different constructs (empty vector EV, *STM2-YFP*, *STM2 E91A-YFP, STM2 E91G-YFP, STM2 E91K-YFP, Strep-STM1*, *Strep-stm1*, *ADR1-L1-Strep*, *STM1-GFP*, *STM1-GFP-NES, STM1-GFP-NLS*, *EDS1-Myc, PAD4-Myc, STM2-Myc, ROQ1-Myc, RPS4-Myc*) were grown overnight. The cells were resuspended in a solution containing 10 mM MgCl_2_, 10 mM MES (pH 5.6) and 150 μM acetosyringone and injected into tobacco (*N. benthamiana*) leaves. Depending on the experiments, wild-type tobacco or the quadruple mutant *epss* ^68^ (deletion of *NbEDS1*, *NbPAD4*, *NbSAG101a*, and *NbSAG101b*) was used. The total amount of bacteria in each system was set at OD600 1.0, comprising OD600 0.2 for bacteria harboring *STM2*, *ROQ1*, *RPS4*, or *ADR1-L1*, OD600 0.7 for bacterium harboring *STM1*, and OD600 0.1 for bacteria *35S::P19* (a viral suppressor of gene silencing), *EDS1*, or *PAD4*. In the experimental groups that lacked one or more bacteria, they were made up to OD600 1.0 with EV-containing bacterium. To examine the effect of metals, 50 µl of MES buffer solution (pH 5.6) containing 10 mM MgCl_2_ together with 50 - 200 µM CdCl_2_, 400 µM ZnCl_2_, or 100 µM CuCl_2_ was injected to the same spots of leaves at 24 h after *Agrobacterium* infection, with injection of the buffer solution containing 10 mM MgCl_2_ serving as a control. Photographs of cell-death phenotypes were taken under UV light at 2-3 d post infection (dpi).

### Bimolecular Fluorescence Complementation (BiFC) Assays

The sequences of *STM1* and *STM2* open reading frame (ORF) were amplified from Col-0 cDNA. The sequences were cloned into the entry vector pDONR and then into the plant expression vectors pGQL-1221-YN (N-terminal half of YFP) and pGQL-1221-YC (C-terminal half of YFP) using Gateway cloning. The constructs were transfected into *Agrobacterium tumefaciens* strain GV3101. After overnight culture, bacterial cells were harvested by centrifugation and resuspended in a buffer (10 mM MES, pH 5.7, 10 mM MgCl_2_, 200 mM acetosyringone) at OD_600_ of 0.2. Equal amounts of a mixture of *Agrobacterium* suspensions carrying N-terminal YFP and C-terminal YFP constructs were injected into 4-week-old tobacco leaves. Samples were taken for viewing or for extracting protoplasts at 2-3 dpi. To test the effect of Cd, 200 µl of MES buffer (pH 5.6) containing 0 or 200 µM Cd was injected 6 h prior to sampling viewing. Protoplasts were isolated from tobacco leaves by incubating leaf segments with an enzyme solution (1% cellulase R-10, 0.25% macerozyme R-10, 0.5 M mannitol, 10 mM CaCl_2_, 20 mM MES-KOH, 20 mM KCl, 0.1% BSA, pH 5. 6) on a shaker at 37°C in the dark for 2-3 h. The enzymatic reaction was terminated by addition of a W5 solution (154 mM NaCl, 125 mM CaCl_2_, 5 mM KCl, 5 mM glucose, 0.03% MES, pH 5.6) and the protoplasts were resuspended in W5. The YFP and chloroplast fluorescence of the isolated protoplast were observed at 525 nm and 640 nm for emission, and at 510 nm and 675 nm for excitation, respectively, under laser confocal microscope (SP8, Leica).

### Blue Native PAGE (BN-PAGE)

*Agrobacterium tumefaciens* strain GV3101 harboring *STM2-YFP*, mutated p-loop motif *STM2-p-loop-YFP* (the conserved p-loop motif favors protein multimerization), *EDS1-Myc*, *PAD4-Myc*, and *ADR1-L1-Strep* constructs were generated and co-expressed in wild-type or *epss* leaves as described in the tobacco HR assays. Four leaf discs (5 mm diameter) were collected at 30 h after infection. To test the effect of Cd, 50 µl of MES buffer (pH 5.6) containing 0 or 200 µM Cd were injected at 24 h after *Agrobacterium* infection and leaf discs were collected at 36 h. Samples were immediately homogenized with liquid nitrogen and extracted with a NativePAGE™ Sample Buffer (BN2008, Invitrogen™) with 1× protease inhibitor cocktail (MCE). The extracts were centrifuged at 20,000 *g* for 30 min at 4°C. Native PAGE™ 5% G-250 (BN2004, Invitrogen™) at the final concentration of 0.125% was added to the supernatant. The supernatant (10 µl) was taken as a sample of non-denatured proteins to be separated by electrophoresis through Native PAGE™ Novex® 3 to 12% Bis-Tris gels (BN1001, Invitrogen™). Electrophoresis was conducted with an anodic buffer (1 l comprising 50 ml 20x NativePAGE Buffer and 950 ml distilled water) in the outside tank and a cathodic Dark Blue buffer (200 ml comprising 10 ml 20x NativePAGE Buffer, 10 ml 0.4% G-250 and 180 ml distilled water) in the inside tank and 150 V was applied for 30 min. The inside tank was switched to a cathodic Light Blue buffer (200 ml comprising 10 ml 20x NativePAGE Buffer, 1 ml 0.4% G-250 and 189 ml distilled water) and 150 V was applied for 90 min. Another 10 μl of the supernatant was taken and added to the loading buffer after boiling and separated by SDS-PAGE. Subsequently, gels were transferred to PVDF membranes and STM2-YFP detected by anti-GFP antibody (1:5000, G1544, Sigma), EDS1-Myc and PAD4-Myc by anti-Myc antibody (1:5000, M4439, Sigma), andADR1-L1-Strep by anti-Strep antibody (1:5000, ab76949, Abcam).

### Expression and purification of recombinant proteins

STM1 was expressed in Sf9 insect cells. STM2 was expressed in Sf9 insect cells for the pull-down experiment and in human HEK293T cells for all other experiments. For expression in insect cells, CDS of *STM1* was cloned into the N-terminal Flag-tagged pFastBac Ⅰ plasmid (Invitrogen), and the construct was transformed into DH10Bac (Invitrogen) to obtain a recombinant bacmid, which was transfected into Sf9 insect cells (Invitrogen) to produce recombinant baculovirus. The recombinant baculovirus was further used to infect Sf9 cells and cultured at 28℃ on a shaker (100 rpm) for 48-60 h before cells were collected. For expression in human HEK293T cells, the Flag sequence was fused to the N-terminus of *STM2* CDS, TIR (1-185 AA) or LRR (550-1261 AA) domain of STM2 sequence. The Strep sequence was fused to the N-terminus of *STM1* CDS by PCR. The fragments were cloned into the pMlink vector with an N-terminal Protein A-SUMO tag. HEK293T cells were transfected with the plasmids construct using PEI reagent (40816ES02, Yeasen). When multiple plasmids were co-expressed, equal amounts were used. The cells were cultured at 37°C and 5% CO_2_ on a shaker (100 rpm) for 3 d. For protein purification, the expressed Sf9 or HEK293T cells were collected by centrifugation at 6500 *g* for 10 min and resuspended in a lysis buffer containing 50 mM Tris-HCl (pH 7.5), 300 mM NaCl, 10% glycerol, 5 mM ATP, 1 mM EDTA, 0.2% Chaps (20102ES08, Yeasen), and 1×Protease Inhibitor Cocktail (HY-K0010, MCE,) at a 1:5 (v/v) ratio of cells to the lysis buffer. The sf9 cells were lysed by high-pressure fragmentation (600 MPa) and HEK293T cells were collected into 50 ml centrifuge tubes, placed on a spinner and spun for 2 h at 4°C to rupture them. The contents were centrifuged at 20,000 *g* for 1 h. The supernatant was transferred to a new tube, to which anti-Flag Affinity Beads (1/200 of the supernatant volume, SA042500, Smart-lifesciences) or IgG Beads (1/50 of the supernatant volume, SA082500, Smart-lifesciences) were added.

After incubation at 4℃ for 2-4 h, the beads were washed three times with a wash buffer (50 mM Tris-HCl, pH 7.5, 300 mM NaCl, 10% glycerol). The washed beads were collected in 1.5 ml microcentrifuge tubes, and initial purified protein was eluted from anti-Flag Affinity Beads with wash buffer supplemented with 3xFLAG peptide (0.5 mg/ml). The IgG beads were incubated with a washing solution containing 2 µM Ulp1 protease at 4°C for 30 min to release the eluted protein with the Protein A-sumo tag removed. The content was centrifuged at 10,000 *g* at 4°C for 2 min. The eluted protein in the supernatant was collected, concentrated using Amicon Ultra-15 or 50 centrifuge filter units (Millipore) and applied onto a Superdex 6 HiLoad 10/300 size-exclusion column (Cytiva) pre-equilibrated with a gel-filtration buffer consisting of 10 mM HEPES (pH 7.5), 150 mM NaCl and 1 mM DTT. Protein fractions were detected by SDS-PAGE with Coomassie brilliant blue staining and immunoblot analysis. The protein-containing fractions were concentrated using Amicon Ultra-15 or 50 centrifuge filter units (Millipore). Protein samples were stored at -80°C before analysis.

### Immunoblotting analysis

For the detection of plant-derived proteins, soluble proteins in *Arabidopsis* transgenic plants or tobacco transient expression leaves were extracted with a solution of 5% SDS,10 mM EDTA and 100 mM NaCl. Samples were denatured by adding 0.2 times the volume of a protein loading buffer (20315ES20, Yeasen), heated in a boiling water bath for 10 min, and then separated on 8% or 10% SDS-PAGE. Subsequently, proteins were transferred onto a polyvinylidene difluoride (PVDF) membrane and probed with antibodies in the TBST solution (100 mM Tris-HCl, 150 mM NaCl, 0.1% (v/v) Tween 20, pH 7.4) with 5% nonfat dry milk. Anti-GFP antibody (1:5000, G1544, Sigma) or an anti-Strep antibody (1:5000, ab76949, Abcam) were used as the primary antibodies and HPR-conjugated goat anti-rabbit/mouse Ig was used as the secondary antibody (IMR-GtxRb-003-DHRPX /GtxRt-003-DHRPX ,1:5000, Jackson). Chemiluminescence was performed using Super ECL Detection Reagent (36208ES60, Yeasen) and Tanon 5200 apparatus (Tanon Technology) exposure blots. Finally, the PVDF membrane was stained in Ponceau S (BL519A, Biosharp) for 30 min, washed twice with deionized water and photographed.

### Co-immunoprecipitation (CoIP)

CoIP assays were performed by transient expression in *N. benthamiana* leaves. The coding sequences of *STM1* and *STM2* were amplified by PCR and cloned into pUC19 entry vectors. Using Gateway™ LR Clonase™ II (11791020, Invitrogen™), *YFP*, *STM1* and *STM2* were recombined into the modified plant expression vector pEarly 101-1 (C-terminal YFP with 35S promoter). *STM1* and *STM2* were also recombined into the modified plant expression vector pEarly 101-3 (N-terminal Strep Ⅱ with 35S promoter). These constructs were transiently expressed in tobacco leaves using *Agrobacterium tumefaciens* strain GV3101. The total protein was extracted from tobacco leaves using a buffer solution (40 mM Tris pH 7.5, 150mM NaCl, 5 mM EDTA, 1% Triton, 5 mM DTT, 1×protease inhibitor cocktail (MCE). The extracts were centrifuged at 20 000 *g* for 1 h at 4°C. 20 µl of supernatant was used as the input and the rest was transferred to a new centrifuge tube, to which 10 μL GFP-Trap agarose beads (gta-20, Chromotek) were added. After incubation at 4 °C for 3 h, the beads were washed five times with a washing buffer (40 mM Tris pH 7.5, 150 mM NaCl, 5 mM EDTA, 0.5% Triton). Protein-bound beads were used as the immunoprecipitated samples. The immunoprecipitated proteins and input proteins were detected by immunoblotting with anti-GFP antibody (1:5000, G1544, Sigma) or anti-Strep antibody (1:5000, ab76949, Abcam).

### Pull-down Assays

The coding sequences of *STM1* and *STM2* were cloned into the pFast Bac Ⅰ vector with a C-terminal GFP or MBP tag. The *STM1-GFP*+*STM1-MBP*, *STM1-GFP*+*STM2-MBP* and *STM2-GFP*+*STM2-MBP* plasmids were co-expressed and transformed into Sf9 cells. Protein expression and purification were performed as described above using GFP-Trap agarose beads (gta-20, Chromotek). 20 µl of lysate from Sf9 cells without the added beads were used as the input. The GFP-Trap agarose beads were washed four times with a washing buffer (40 mM Tris, pH 7.5, 150 mM NaCl, 5 mM EDTA, 0.5% Triton) to remove unbound proteins. Protein-bound beads were used as the pull-down samples. The pull-down and input proteins were separated by SDS-PAGE and visualized by western blot using anti-GFP antibody (1:5000, G1544, Sigma) or anti-MBP antibody (1:5000, SAB2104172, Sigma).

### NADase Assays

To perform *in vivo* NADase assay in *E. coli*, the coding sequences of STM1-TIR (1-165 AA), STM2-TIR (9-181 AA), and RPS4-TIR (1-183 AA) were cloned into the pET-29a vector and transfected into *E. coli* BL21 strain. Cell cultures were grown at 37 °C to OD_600_ of 0.6. Expression of proteins was induced for 2 h by the addition of 0.3 mM isopropyl β-D-thiogalactoside. Cells were washed with distilled water and resuspended to OD600 of 0.5. One ml of the bacterial solution was taken and centrifuged. 150 µl of 1 M HClO_4_ was added and the samples were placed on ice for 10 min. After centrifugation at 4°C for 10 min, the enzymatic reaction was stopped by the addition of 50 µl of 3 M K_2_CO_3_. The content was centrifuged at 12,000 *g* at 4 °C for 10 min. The supernatant was stored at -80 °C prior to HPLC analysis. NAD^+^ concentration in the supernatant was determined as described below.

To determine NADase activity *in vitro*, the purified recombinant proteins of STM1 and STM2 were resuspended on anti-Flag Affinity Beads. The concentrations of proteins on beads were determined by the Bradford method and were diluted 50 μM. Ten µl of anti-Flag Affinity beads with purified protein were incubated with 40 μl 30 μM NAD^+^ (final concentration) in a reaction buffer (92.4 mM NaCl and 0.64x Phosphate Buffer Saline, containing 6.4 mM phosphate). Reactions were conducted at room temperature (25°C) for 3 h and stopped by the addition of 50 µl of 1 M HClO_4_. The samples were placed on ice for 10 min. The acid in the sample was neutralized by the addition of 16.7 µl of 3 M K_2_CO_3_, placed on ice for 10 min and then separated by centrifugation. In some experiments, Cd, Zn or Cu were added to the reaction at a final concentration of 50 µM, with no metal addition as the control. The extracts were stored at -80 °C prior to LC-MS/MS analysis.

*STM1* and *STM2* constructs were transiently expressed in tobacco leaves using *Agrobacterium tumefaciens* GV3101. Leaf samples were collected at 26 or 36 h post infiltration for NADase metabolite assays. To test the effect of Cd, 50 µl of MES buffer (pH 5.6) containing 0 or 200 µM Cd was injected at 24 h after infiltration of *Agrobacterium* and leaf samples were collected at 36 h after infiltration. Four leaf disks (8 mm in diameter) from 3 different plants were snap-frozen in liquid nitrogen and homogenized into powder. The sample was dissolved in 200 µl 50% (v/v) methanol and vortexed for 1 h. Plant extracts were centrifuged at 13 000 *g* for 15 min. The supernatant was transferred and stored at -80°C prior to LC-MS/MS analysis.

### Determination of NAD^+^ and NAD^+^ metabolites by HPLC or LC-MS/MS

The concentration of NAD^+^ in *E. coli* was determined by HPLC using a LC-18-T column. The mobile phase consisted of buffer A (0.05 M Phosphate Buffer, pH 6.0) and buffer B (100% methanol) with the following elution gradient, 0 - 5.00 min, 100% A; 5.00 - 6.00 min, 100% to 95% A; 11 - 13 min, 95% to 85% A; 23 - 24 min, 85% to 100% A. The flow rate was 1 ml min^−1^ and the column temperature was maintained at 40°C. The NAD^+^ peak appeared at 11 min. The UV absorption of NAD^+^ was detected at 260 nm. A calibration curve was established between the peak area of NAD^+^ and the concentration of the NAD^+^ standard.

In tobacco leaf extracts and in vitro NADase assays, NAD^+^ and NAD^+^ metabolites were determined by LC-MS/MS. Tobacco leaf extracts or protein reaction samples (100 µl) were mixed with 100 µl 50% methanol and centrifuged at 12,000 *g* for 10 min. The supernatant was separated with an ACQUITY UPLC HSS T3 column (Waters). The mobile phase consisted of 2 mM ammonium acetate in water (A) and 100% methanol (B), using the following elution gradient, 0 - 2.00 min, 1% B; 2.00 - 7.00 min, 1% to 15% B; 7.00 -9.00 min, 15% to 95% B; 9.00 - 9.10 min, 95% to 1% B; 9.10 - 12.00 min, 1% B. The flow rate was 0.3 ml min^−1^ and the column temperature was maintained at 40°C. NAD^+^ metabolites were detected and quantified with a Triple Quad mass spectrometer (6500; Agilent). Multiple Reaction Monitoring conditions were optimized using authentic standard chemicals including NAD^+^ ([M + H] + 664 > 136.00, 664 > 428, 664 > 542); Nam ([M + H] + 123 > 80); cADPR ([M + H] + 542 > 136); v-cADPR ([M + H] + 542 >136). The concentrations of metabolites were quantified by using external calibration curves for NAD^+^, ADP-ribose (ADPR) and Nicotinamide (Nam) dissolved in 50% methanol.

### Microscale thermophoresis analysis (MST)

Recombinant STM1 and STM2 proteins, STM2 TIR or LRR domains, and mutated STM2 in which C595, C599, C1109, and H1022 were replaced with Ala were purified as described above. All recombinant proteins contained a Flag tag. The proteins (10 µM) were labelled with a fluorescence dye (NT-647-NHS dye) as the assay target proteins according to the method provided in the Monolith^TM^ RED-NHS Second Generation Protein Labelling Kit (MONTEMPER, MO-L011). In the metal binding assay, the final concentration of the target protein was 20 nM. The ligand solution of CdCl_2_, ZnCl_2_, or CuCl_2_ was diluted by the MST buffer gradient. Affinity assays were performed on a Monolith NT.115 instrument (Nano Temper, Germany) with a Pico-Red detector, using automatic detection of LED power (20-90%) and MST power (medium or high). Data were evaluated using the MO affinity analysis software.

### Inoculation with *Ralstonia solanacearum* and disease assessments

*R. solanacearum* strains GMI1000 and RS1115 were grown in Blue-Green Medium (BG) for 36 h at 28°C. The bacterial cultures were centrifuged at 8000 *g* for 10 min and resuspended in sterile water to OD_600_ of 0.55. *Arabidopsis* plants were grown in puffy Jiffy-7 peat pellet (Jiffy Products, Norway, containing 43.0% organic matter) for 4 weeks and the growth medium was then infiltrated with *R. solanacearum* by soaking 50 Jiffy-7 pellets in a tray with 2 L of the bacterial suspension for 30 min. The pellets were then transferred to new trays. The degree of plant wilt was quantified according to the method described previously with the disease score ranging from 0 for healthy plants to 4 for dead plants^69^. Plants were monitored for wilting symptoms from 4 to 16 days after infiltration. To investigate the effect of Cd or Cu on *R. solanacearum* infection, Jiffy-7 pellets were soaked with 1 mM Cd or Cu solution or distilled water for 3 h and *R. solanacearum* was inoculated 48 h later.

### Determination of metal concentrations

Roots and shoots of plants grown in agar plates or Jiffy-7 peat pellets were collected, washed with deionized water, blotted dry, and frozen in liquid nitrogen. Cell saps were obtained by centrifugation at 20,000 *g* at room temperature. In the soil experiment, available metals in the soil were extracted with 0.01 M CaCl_2_ (1:5, w/v). The concentrations of metals in the cell saps and the soil extracts were determined by using inductively coupled plasma mass spectrometry (ICP-MS, Perkin-Elmer NexION 300x).

### Statistical analysis

All experiments were repeated at least twice with similar results. The data are presented as the mean ± SD. All data were analyzed using Microsoft Office 365 (Excel) and GraphPad Prism 8.0. Significant differences between two sets of data were determined by two-sided student’s *t-*tests. The data from the experiments of genotype x Cd or Cu x inoculation of *R. solanacearum* involving more than one factor were analyzed by analysis of variance (ANOVA) assuming a Gaussian distribution, followed by Tukey’s post-hoc multiple comparisons. The probability levels (ns, not significant, \**P* < 0.05, \*\**P* < 0.01 and \*\*\**P* < 0.001) are indicated in the Figures. Sample sizes are indicated in the Figure legends. Diagrams were drawn using GraphPad Prism8.0, or “ggplot2” and “pheatmap” in R packages.

## Acknowledgments

We thank Dr. Yanfei Mao for providing the single base editing vectors (CBEmax-nCas9NG-p19 and ABE max-nCas9NG-p19) and Dr. Gaofei Jiang for providing *R. solanacearum* strains GMI1000 and RS1115.

## Funding

This work was supported by the National Natural Science Foundation of China (No. 42090062). Li Wan was supported by Key Laboratory of Plant Design, CAS Center for Excellence in Molecular Plant Sciences, Institute of Plant Physiology and Ecology, Chinese Academy of Sciences and National Natural Science Foundation of China project 32270304. Jeffery L. Dangl was supported by the National Science Foundation grant IOS-1758400 and Howard Hughes Medical Institute.

## Author contributions

F.J.Z. and L.W. designed the research and supervised the project. C.G., S.S.C., and J.C. performed the experiments. C.G. and T.Z. performed data analysis and visualization. C.G., X.Y.H., P.W. S.M.D., J.D., L.W. and F.J.Z. discussed the data. C.G. and F.J.Z. performed the writing - original draft. F.J.Z., L.W. and J.D. performed the writing – review & editing.

## Declaration of interests

The authors declare no competing interests.

## Data availability

The RNA-seq data are deposited in NCBI GEO database (GSE291114 for *cad1-3* and *stm1-1 cad1-3*; GSE291226 for Col-0, *stm1-2*, *stm2-2, stm2-2 stm1-2*).

## Code availability

No customized code was generated in this study.

---

**Extended Data Fig. 1:**
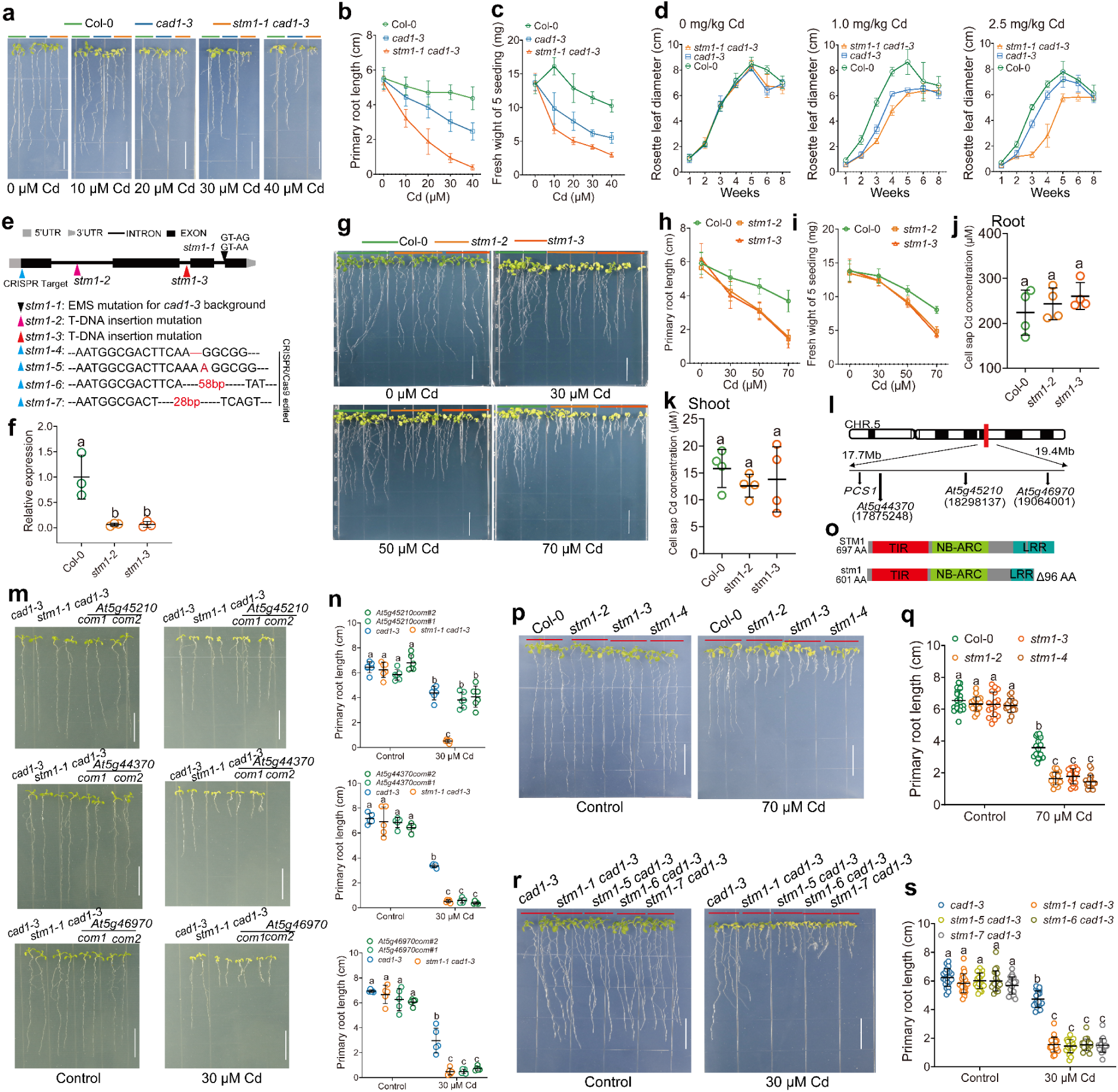
Cd-sensitive phenotype of *stm1* or *stm1-1 cad1-3* mutants and identification of the causal gene. **a-c** Growth phenotype (**a**), primary root length (**b**) and fresh weight of 5 plants (**c**) of Col-0, *cad1-3*, and *stm1-1 cad1-3* grown on agar plates containing different concentrations of Cd. **d** Effect of Cd on growth (measured as rosette leaf diameter) of Col-0, *cad1-3*, *stm1-1 cad1-3* grown in soil amended with 0, 1.0 or 2.5 mg kg^−1^ of Cd. **e, f** Information of various *stm1* mutants used in the study. (**e**) Sketch of different *stm1* mutant alleles in the gene structure of *STM1* (top); basic information for *stm1-1* to *stm1-7* (bottom). (**f**) Expression of *STM1* in the roots of *stm1-2* and *stm1-3* (T-DNA insertion mutants) relative to Col-0. **g-i** Growth phenotype (**g**), primary root length (**h**) and fresh weight of 5 plants (**i**) of Col-0, *stm1-2*, and *stm1-3* grown on agar plates containing different concentrations of Cd. (**j, k**) Cd concentration in the cell sap of roots (**j**) and shoots (**k**) of Col-0, *stm1-2*, and *stm1-3* grown on agar plates for 10 days and then exposed to Cd (70 µM) for 24 h. **l** Mapping of candidate genes by bulked segregant analysis combined with whole-genome sequencing. **m, n** Complementation of *AT5G45210*, but not *At5g44370* or *At5g46970*, rescued the Cd sensitivity phenotype of *stm1-1 cad1-3*. Growth phenotype (**m**) and primary root length (**n**) of various genotypes grown under control or 30 µM Cd. Two independent complementation lines were used for each candidate gene.**o** stm1 from *stm1-1 cad1-3* cDNA is truncated missing the last 96 amino acids compared to wild-type STM1. **p, q** Knockout of *STM1* in Col-0 leads to Cd sensitivity; growth phenotype (**p**) and primary root length (**q**). **r, s** Knockout of *STM1* in the *cad1-3* background enhanced Cd sensitivity; growth phenotype (**r**) and primary root length (**s**). Data are means ± SD with n = 20 individual plants in (**b**); n = 4 biological replicates in (**c**); n = 3 biological replicates in (**d**); n = 30 individual plants in (**h**); n = 6 biological replicates in (**i**). All data points are shown with means ± SD, n = 3 biological replicates in (**f**); n = 4 biological replicates in (**j, k**); n = 5 individual plants in (**n**); n = 17-18 individual plants in (**q**); n = 20-21 individual plants in (**s**). Each replicate comprising 3 plates of >100 seedlings in (**f, j, k**).In (**f, j, k**) and (**n, q, s**), different letters represent significance at *P<*0.05 (one- and two-way ANOVA, respectively, followed by Tukey’s post-hoc multiple comparisons). Scale bars are 1.5 cm in (**a, g, m, p, r**).

**Extended Data Fig. 2:**
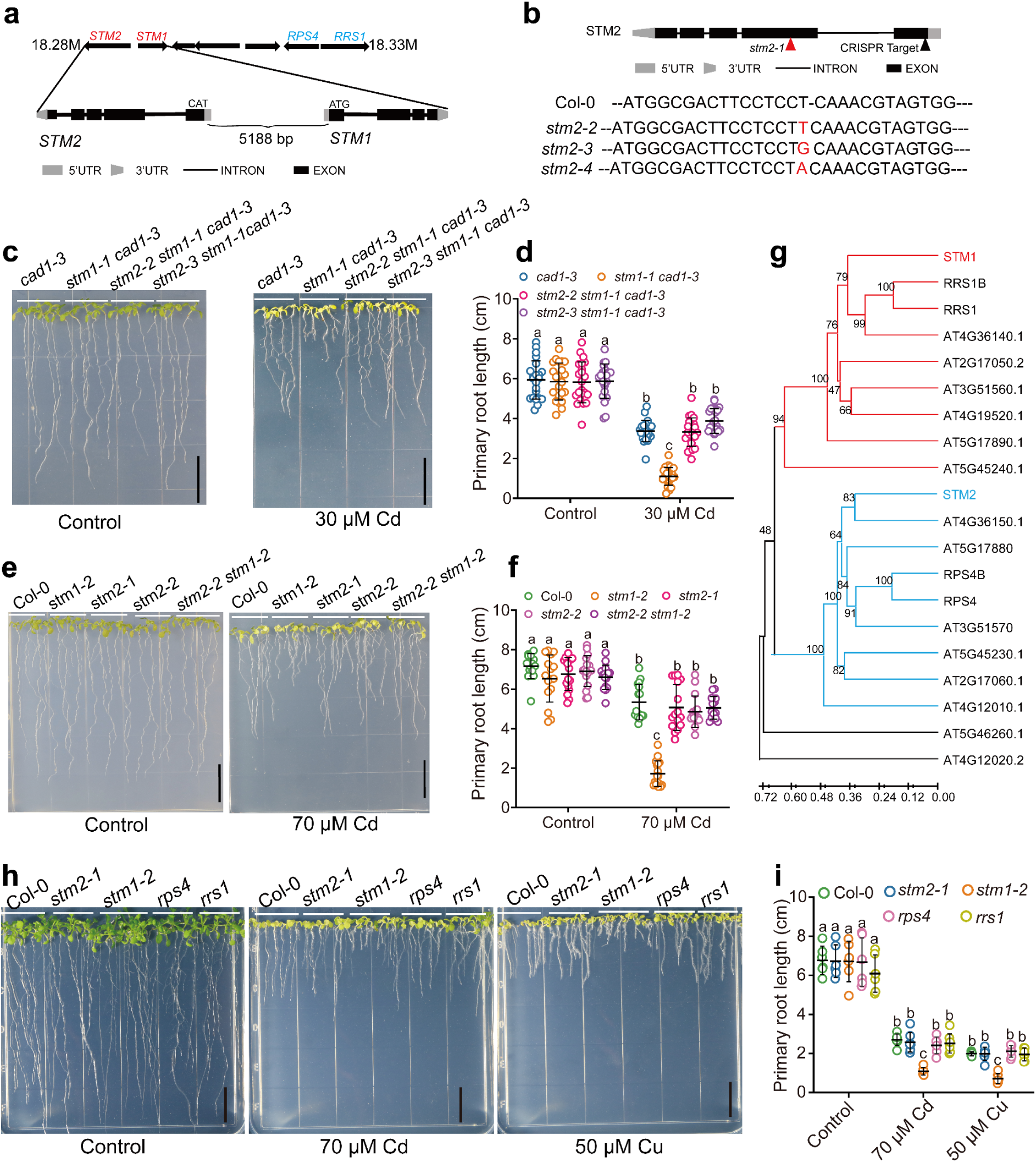
Knockout of *STM2* rescues the Cd-sensitive phenotype of *stm1-2 or* stm1-1 cad1-3. **a** Schematic gene structure of *STM1* and *STM2* in a NLR gene cluster. **b** Sketch of the *STM2* gene structure and genotypes of *STM2* knockout lines by CRISPR/Cas9; red and black triangles indicate the T-DNA insertion and CRSIPR/Cas9 editing target, respectively. **c, d** Cd sensitivity in *stm1-1 cad1-3* was rescued by knockout of *STM2*; growth phenotype (**c**) and primary root length (**d**). **e, f** Knockout of *STM2* in Col-0 background did not affect Cd sensitivity, but knockout of *STM2* in *stm1-2* background rescued its Cd sensitive phenotype; growth phenotype (**e**) and primary root length (**f**). *stm2-1* is a T-DNA insertion line in the Col-0 background. **g** Phylogenetic analysis of STM1 and STM2 with other NLR pairs. STM1 and STM2 are clustered in two separate groups and fall among the head-to-head TNL pairs. **h, i** Knockout of *RPS4* or *RRS1* did not affect Cd or Cu tolerance in *Arabidopsis thaliana*; growth phenotype (**h**) and primary root length (**i**) of Col-0, *stm1-2, stm2-1, rps4 and rrs1* grown on agar plates with or without Cd or Cu. All data points are shown with means ± SD, n = 22 or 23 individual plants in (**d**), n = 15-17 individual plants in (**f**), and n = 6-7 individual plants in (**i**). In (**d, f, i**), different letters represent significance at *P<*0.05 (two-way ANOVA, followed by Tukey’s post-hoc multiple comparisons). Scale bars are 1.5 cm in (**c, e, h**).

**Extended Data Fig. 3:**
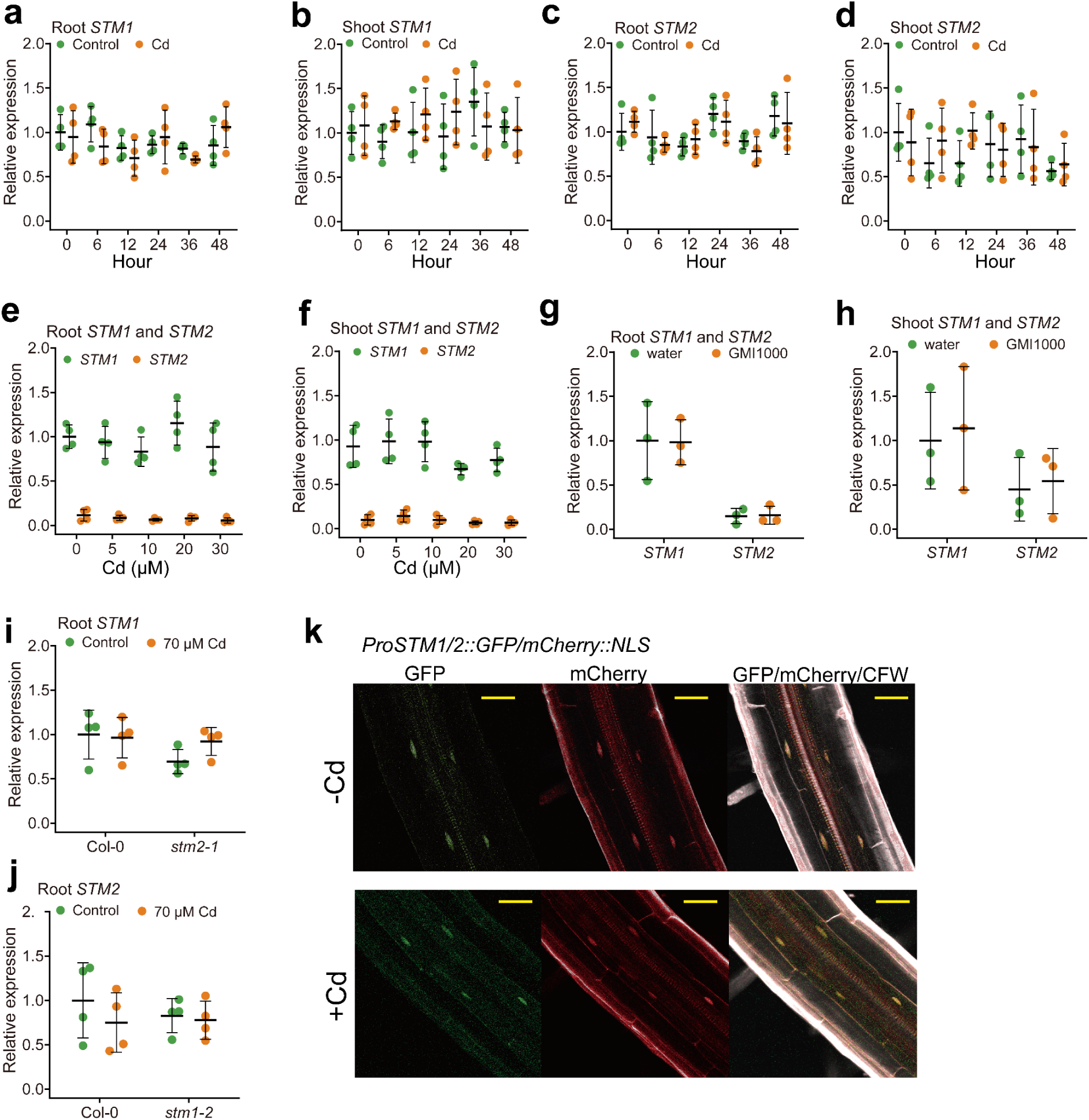
Cd and *Ralstonia solanacearum* infection do not alter the expression of *STM1* and *STM2*. **a-d** Expression of *STM1* (**a, b**) and *STM2* (**c, d**) in the roots (**a, c**) and shoots (**b, d**) of Col-0 plants exposed to 70 µM Cd for 0 – 48 h. **e, f** Expression of *STM1* and *STM2* in roots (**e**) and shoots (**f**) of *cad1-3* plants exposed to 0 – 30 µM Cd for 12 h. **g, h** Expression of *STM1* and *STM2* in the roots (**g**) and shoots (**h**) of Col-0 plants infected with or without *Ralstonia solanacearum* (GMI1000). Plants were grown in Jiffy pellets for 4 weeks and inoculated with water or GMI1000 for 24 h before sampling of roots and shoots. **i** Expression of *STM1* in Col-0 and *stm2-1* plants treated with or without Cd. **j** Expression of *STM2* in Col-0 and *stm1-2* plants treated with or without Cd.Gene expression levels are relative Col-0 or *cad1-3* in the control with *UBQ10* as an internal reference. All data points are shown in (**a-j**) with means± SD; n = 4 in (**a-f, i, j**) and n = 3 in (**g, h**). **k** Exposure to Cd (70 µM) for 24 h did not affect the spatial expression pattern of STM1-GFP or STM2-mCherry in the roots of transgenic plants expressing *ProSTM1/2::STM1-NLS-GFP/STM2-NLS-mCherry* plants (see Fig. 2f for the construct). Yellow bar = 50 µm.

**Extended Data Fig. 4:**
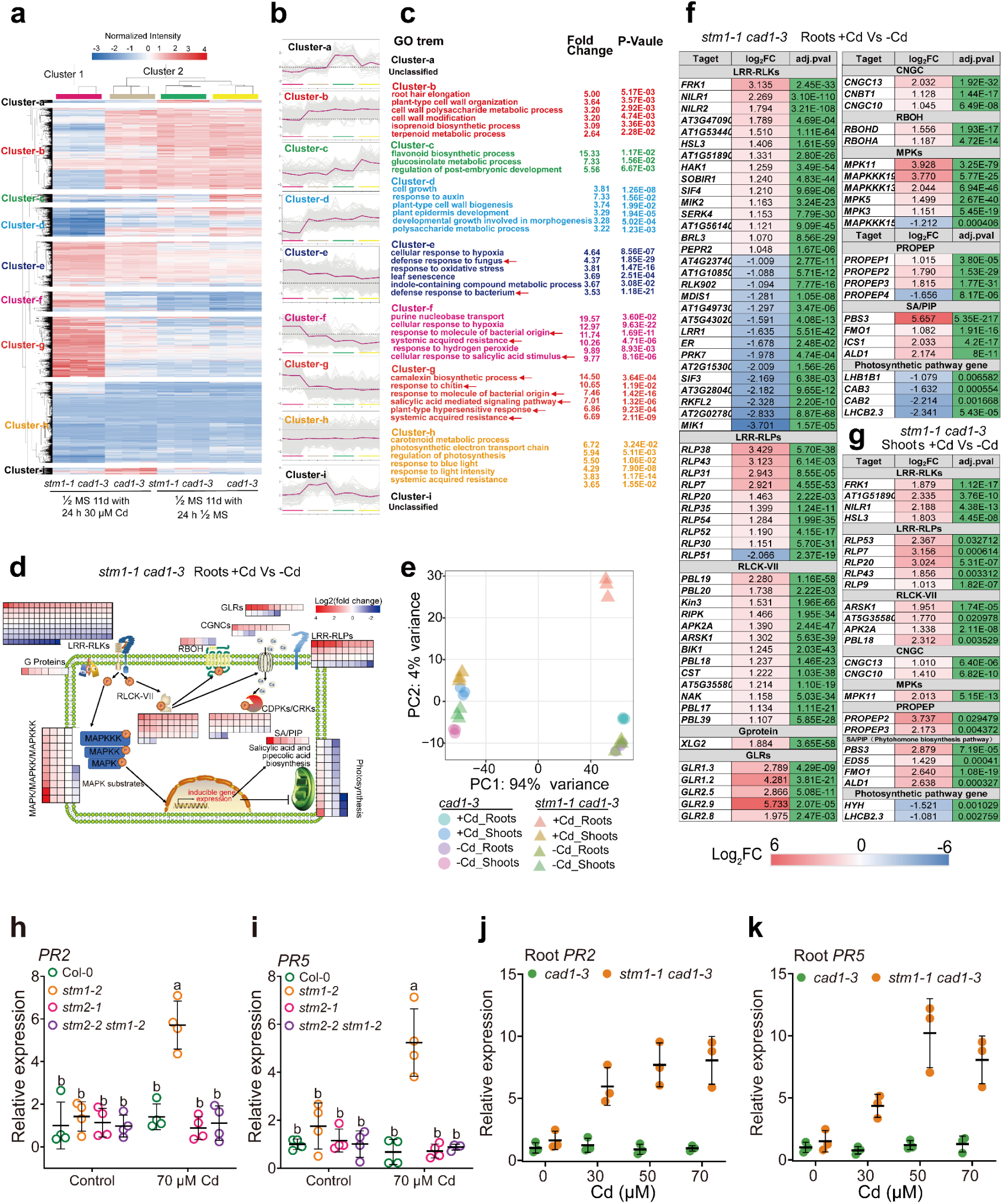
Cd treatment upregulates immunity-related genes in the roots of *stm1 cad1-3*. **a-c** RNA-seq reveals differentially expressed genes (DEGs) between the roots of *cad1-3* and *stm1-1 cad1-3* treated with or without 30 µM Cd for 24 h. Gene expression heatmap showing that Cd caused large transcriptional changes in the Cluster 1 genes in the *stm1-1 cad1-3* roots (**a**). The gene expression patterns were further clustered into 9 clusters (Cluster a to Cluster i), and each cluster was analyzed for expression trends (**b**) and GO enrichment (**c**). Red arrows indicate GO terms associated with plant immunity. **d** Cd-triggered immune responses in *stm1-1 cad1- 3* are similar to ETI^AvrRPS4^. Heatmap represents log_2_-transformed fold changes in gene expression of PTI signaling pathway components in *stm1-1 cad1-3* roots after Cd treatment. Red and blue colors indicate up- and down-regulated genes, respectively. PTI signaling pathway components refer to previous report on ETI^AvrRPS4^(*32*). **e** Principal Component Analysis based on gene expression in roots and shoots samples of *cad1-3* and *stm1-1 cad1-3* treated with or without 30 µM Cd for 24 h. **f, g** PTI-related genes with more than two-fold expression differences in roots (**f**) or shoots (**g**) after Cd treatment. Red and blue colors indicate up- and down-regulated genes, respectively. **h-i** Cd induced expression of *PR2* (**h**) and *PR5* (**i**) in *stm1-2* roots, which was abolished by deletion of *STM2* in *stm2-2 stm1-2.* Data are transcript levels relative to Col-0 in the control treatment with *UBQ10* as an internal reference. All data points are shown with means±SD; n = 4. **j, k** Cd (30 – 70 µM) induced expression of *PR2* (**j**) and *PR5* (**k**) in the roots of *stm1-1 cad1-3*, but not in *cad1-3*. Expression levels were relative *cad1-3* in the control with *UBQ10* as an internal reference. All data points are shown with means±SD; n = 3.

**Extended Data Fig. 5:**
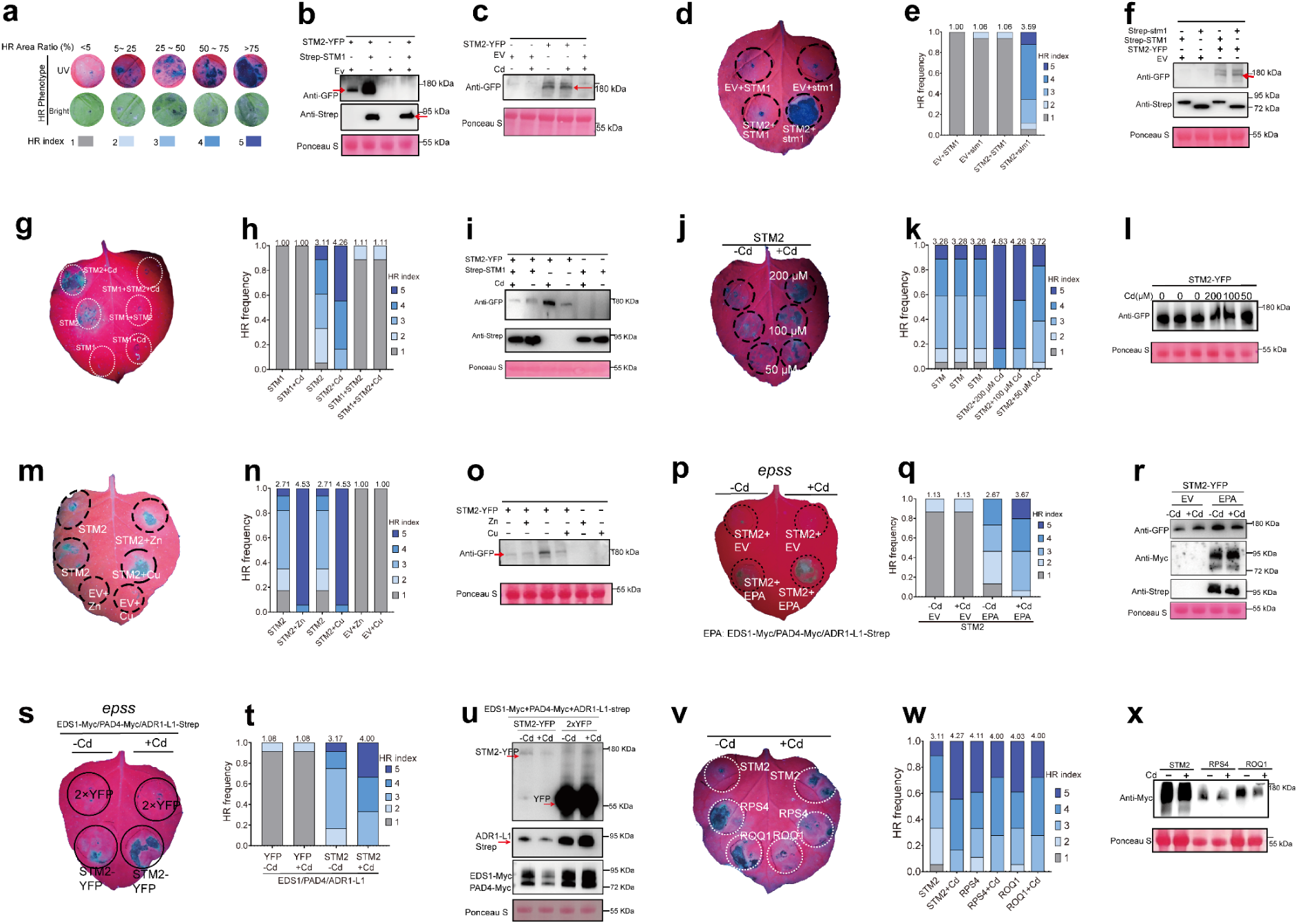
STM2 activated HR in tobacco leaves is enhanced by divalent transition metals and suppressed by STM1. **a** Semi-quantitative scoring of HR based on the proportion of regions producing cell death. All HR test scores are referenced to this scheme. **b** Detection of STM2-YFP and Strep-STM1 fusion proteins expressed in tobacco HR assays shown in Figure 3A using GFP and Strep antibodies, respectively. **c** Detection of STM2-YFP fusion proteins expressed in tobacco HR assays shown in Figure 3B using GFP antibody. **d-f** Mutated *stm1* (truncated *STM1* cDNA from *stm1 cad1-3*) did not inhibit HR caused by STM2. *STM2* with *STM1* or *stm1* were transiently expressed in *N. benthamiana* leaves. Phenotyping under UV light (**d**) and frequency of HR score, n=17 leaves (**e**); detection of STM2-YFP and Strep-STM1 fusion proteins expressed in tobacco leaves by using GFP and Strep antibodies, respectively (**f**). **g-i** STM1 inhibited STM2-induced HR with or without Cd. Transient expression of *STM2*, *STM1* and *STM2 + STM1* in *N. benthamiana* leaves, followed by injection of Cd (200 µM) or buffer solution 24 h later. Phenotyping under UV light (**g**) and frequency of HR score, n=18 (**h**); detection of STM2-YFP and Strep-STM1 fusion proteins expressed in tobacco leaves by using GFP and Strep antibodies, respectively (**i**). **j-l** Dose effect of Cd on STM2-activiated HR in *N. benthamiana* leaves; phenotyping under UV light (**j**) and frequency of HR score, n=18 leaves (**k**); detection of STM2-YFP fusion protein expressed in tobacco leaves by GFP antibody (**l**). **m-o** Effect of Cu (100 µM) or Zn (400 µM) on STM2-activated HR in *N. benthamiana* leaves; phenotyping under UV light (**m**) and frequency of HR score, n=17 leaves (**n**); detection of STM2-YFP fusion protein expressed in tobacco leaves by GFP antibody (**o**). **p-r** Expression of *STM2* in *N. benthamiana epss* quadruple mutant lacking the native *NbEDS1*/*NbPAD4*/*NbSAG101a*/*NbSAG101b* did not cause HR, whereas co-expression of *AtEDS1-AtPAD4-AtADR1-L1* (EPA) restored HR by STM2. Transient expression of *STM2* + *EV* and *STM2* + EPA in *N. benthamiana epss* leaves, followed by injection of Cd (200 µM) or buffer solution 24 h later. Phenotyping under UV light (**p**) and frequency of HR score, n=15 leaves (**q**); detection of STM2-YFP, EDS1-Myc, PAD4-Myc, and ADR1-L1-Strep fusion proteins expressed in tobacco leaves by using GFP, Myc, and Strep antibodies, respectively (**r**). **s-u** Enhancement of HR by Cd requires the presence of STM2. Transient expression of *STM2* or *YFP* with *AtEDS1-AtPAD4-AtADR1-L1* in *N. benthamiana epss* leaves, followed by injection of Cd (200 µM) or buffer solution 24 h later. Phenotyping under UV light (**s**) and frequency of HR score, n=12 leaves (**t**); detection of STM2-YFP, EDS1-Myc, PAD4-Myc, and ADR1-L1-Strep fusion proteins expressed in tobacco leaves by using GFP, Myc, and Strep antibodies, respectively (**u**). **v-x** Cd did not promote HR activated by RPS4 and ROQ1. Transient expression of *STM2*, *RPS4*, or *ROQ1* in *N. benthamiana* leaves, followed by injection of Cd (200 µM) or buffer solution 24 h later. Phenotyping under UV light (**v**) and frequency of HR score, n=18 leaves (**w**); detection of STM2-Myc, RPS4-Myc, and ROQ1-Myc fusion protein expressed in *N. benthamiana* leaves by Myc antibody (**x**). The numbers above the bars in (**e, h, k, n, q, t** and **w**) are the weighted means of HR scores.

**Extended Data Fig. 6:**
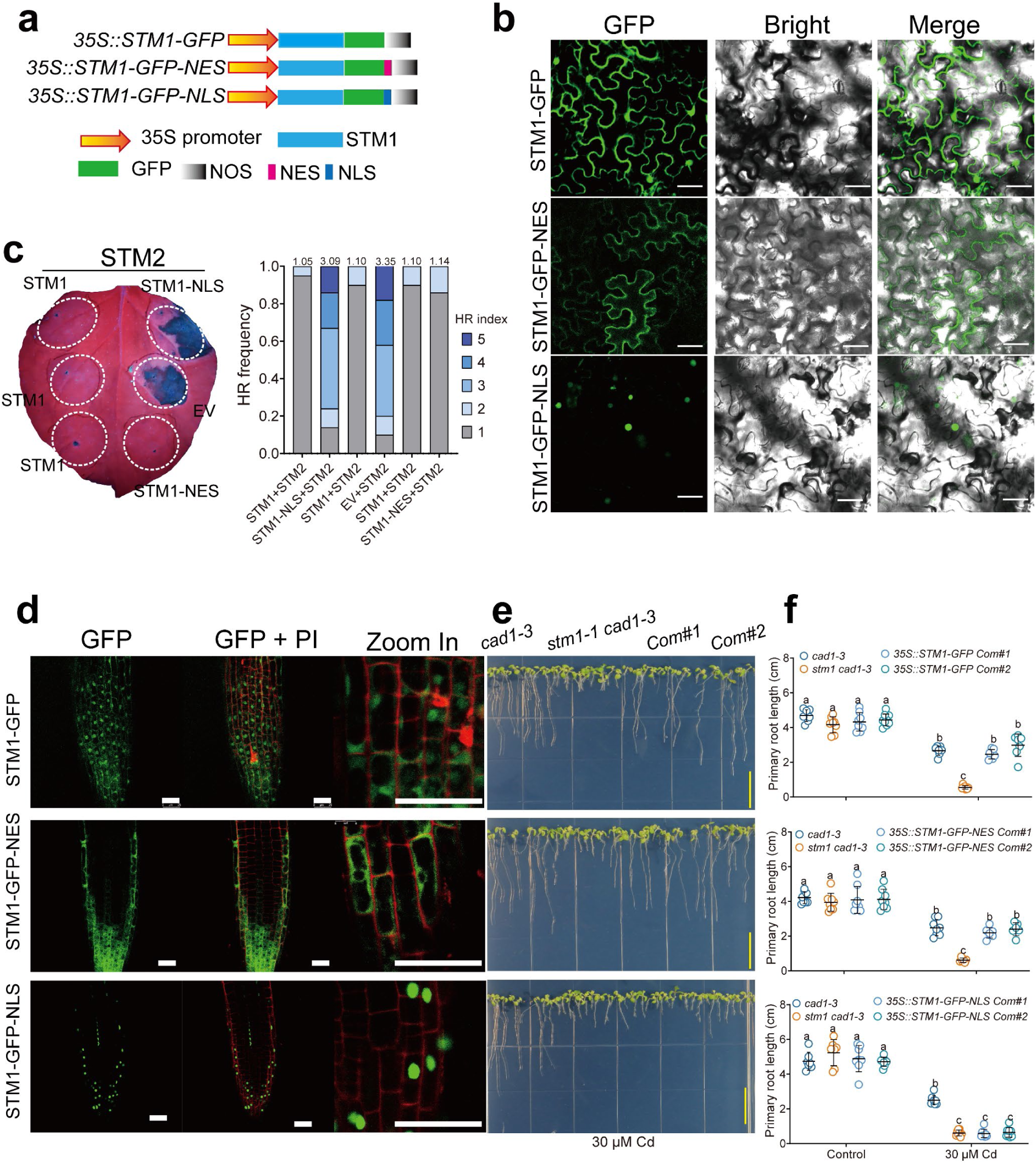
Cytoplasmic localization of STM1 is required for its function. **a** Constructs of *STM1-GFP* with a nuclear export signal (NES) or a nuclear localization signal (NLS) peptide. **b** GFP fluorescence indicates the subcellular localization of STM1-GFP with or without NES or NLS peptide, after transient expression of in epidermic cells of tobacco leaves. White bar = 1.5 cm. **c** HR assay showing the cell death phenotype when different *STM1* constructs listed in (**a**) were co-expressed with *STM2* in tobacco leaves; photographed under UV light (left) and frequency of HR score (right), n=17. The numbers over the bars are weighted means of HR scores. **d-f** Cytoplasmic localization of STM1 is required for Cd tolerance. The different *STM1* constructs listed in (**a**) were complemented to *stm1-1 cad1-3*. GFP fluorescence in the root tips of transgenic plants (**d**). Green, GFP fluorescence; red, PI stain fluorescence. Scale bar =50 µm. Growth phenotype (**e**) and primary root length (**f**) of *cad1-3*, *stm1-1 cad1-3* and transgenic complementation lines grown on agar plates with or without 30 µM Cd. Scale bar = 1.5 cm in (**e**). All data points are shown in (**f**) with means ± SD; n = 7-8 individual plants. Different letters represent significance at *P<*0.05 (two-way ANOVA, followed by Tukey’s post-hoc multiple comparisons).

**Extended Data Fig. 7:**
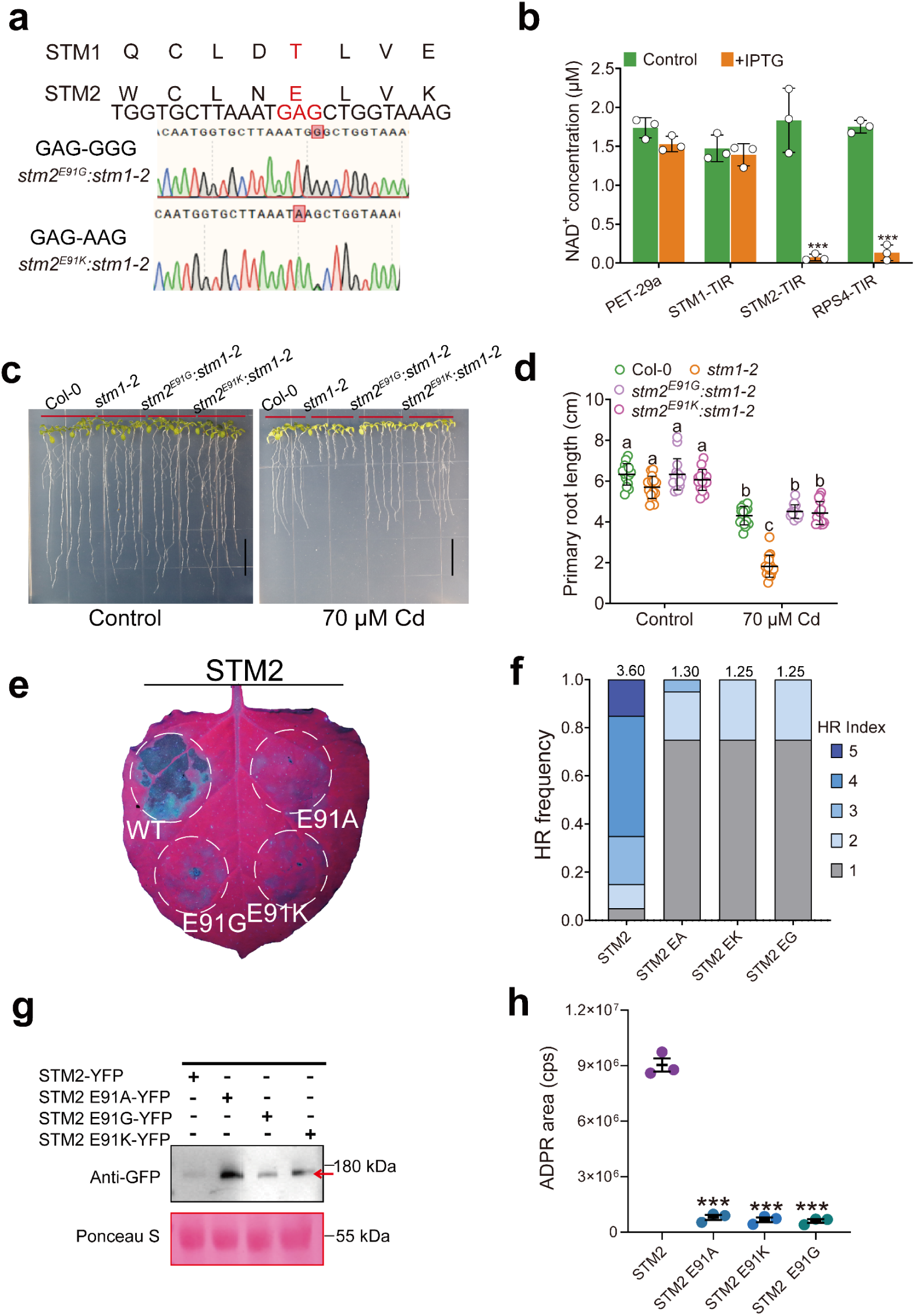
The conserved glutamate residue (E91) in STM2 is necessary for STM2-activiated immune responses in tobacco and Cd sensitivity in *Arabidopsis stm1* mutant. **a** A conserved glutamate residue is present in the TIR domain of STM2 (E91), but not in STM1. Single-base edited lines with E91G and E91K in STM2 in the *stm1-2* background. **b** The NADase activities of the TIR domains of STM1, STM2 and RPS4 expressed in *E. coli*, measured as the depletion of NAD^+^ concentration in the cell lysates after 2 h of induction with or without isopropyl β-D-1 thiogalactopyranoside (IPTG). All data points are shown with means ± SD; n = 3. *** indicates significant difference between +/- IPTG treatments at *P*<0.001 by two-sided *t*-test. **c-d** Single-base editing of *STM2* to replace the conserved glutamate with glycine (E91G) or lysine (E91K) rescued the Cd-sensitive phenotype in *stm1-2*; Growth phenotype (**c**) and primary root length (**d**) of Col-0, *stm1-2*, *stm2 ^E91G^ stm1-2* and *stm2 ^E91K^ stm1-2* grown on agar plates with or without 70 µM Cd. Scale bar = 1.5 cm in (**c**). All data points are shown in (**d**) with means ± SD, n=15. Different letters indicate significant difference at *P*<0.05 (two-way ANOVA, followed by Tukey’s post-hoc multiple comparisons). **e-g** HR assays of STM2, STM2^E91A^, STM2^E91G^ and STM2^E91K^ transiently expressed in *N. benthamiana* leaves; the cell death phenotype under UV light at 3 DPI (**e**); the frequency of HR score with weighted means above the bars, n=20 leaves (**f**); detection of STM2-YFP and mutated STM2-YFP fusion protein expressed in tobacco leaves by GFP antibody (**g**). **h** In vivo NAD^+^ hydrolytic activity of STM2, STM2^E91A^, STM2^E91G^ and STM2^E91K^ transiently expressed in *N. benthamiana* leaves, measured by the production of ADPR in leaf lysates. All data points are shown with means ± SD; n = 3. *** indicates significant difference from STM2 at *P*<0.001 by two-sided *t*-test.

**Extended Data Fig. 8:**
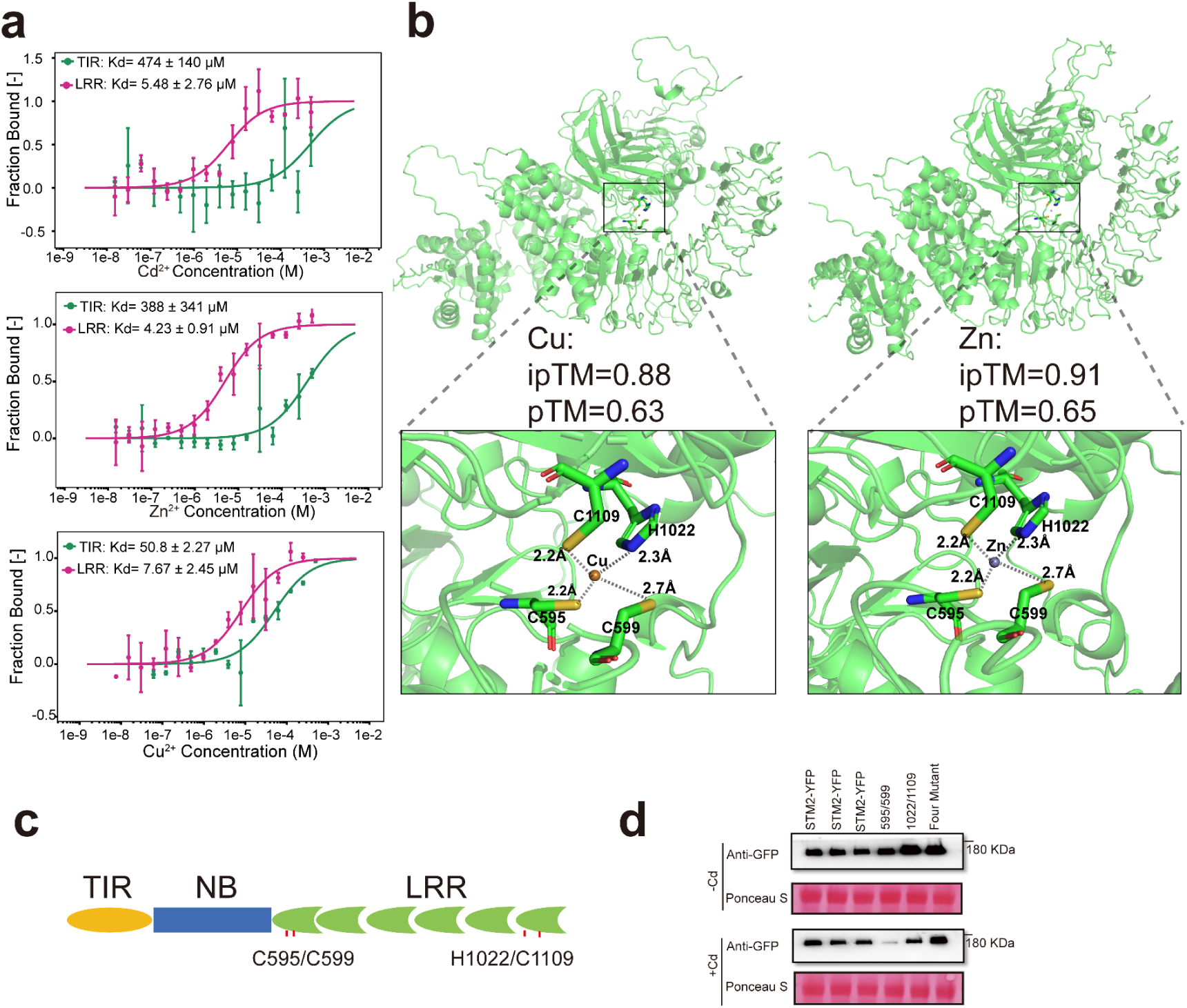
Divalent transition metals are bound to the LRR domain of STM2. **a** MST assays of the binding affinities of the TIR (1-185 AA) and LRR (550-1261 AA) domains of STM2 for Cd^2+^, Zn^2+^, and Cu^2+^. **b** AlphaFold 3 predicts the binding site and the amino acid residues of STM2 for Cu^2+^ and Zn^2+^. The top panel represents the STM2 with Cu or Zn binding and the bottom panels the enlarged view of the white box in the top panel. The metal ion is predicted to be within 3Å distance from the amino acid residues C595, C599, H1022, and C1109. The predicted results are highly plausible (ipTM + pTM > 0.95). **c** Sketch of STM2 with the locations of the predicted binding sites (C595, C599, H1022, and C1109) for transition metals. **d** Detection of STM2-YFP and mutated STM2-YFP fusion protein expressed in *N. benthamiana* leaves in the HR assays shown in Fig. 5g by GFP antibody.

**Extended Data Fig. 9:**
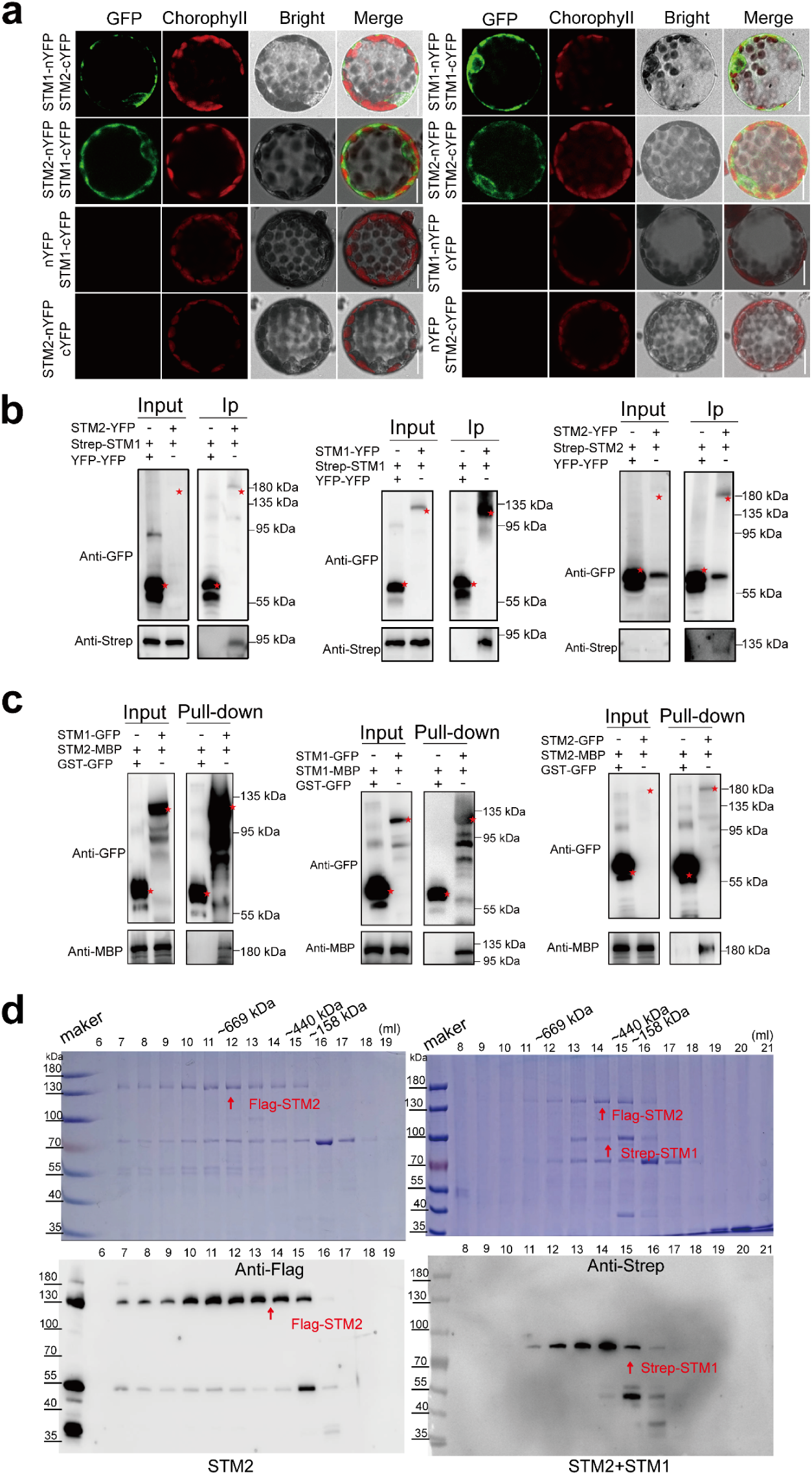
Homomeric and heteromeric interactions between STM1 and STM2. **a** BiFC assays in tobacco leaf protoplasts show that STM1 and STM2 interact with each other (left panels) and between themselves (right panels). The bottom two rows are the negative control. Scale bars = 25 µm. **b** Co-immunoprecipitation assays show that STM1 and STM2 interact with each other (left panels) and between themselves (middle and right panels for STM1 and STM2, respectively). The red star marks the position of the target protein. **c** Pull- down assays show that STM1 and STM2 interact with each other (left panels) and between themselves (middle and right panels for STM1 and STM2, respectively). The red star marks the position of the target protein. **d** Gel filtration of Flag-STM2 alone (Left panels) and the mixture of Flag-STM2 and Strep-STM1 (right panels) recombinant proteins. Eluate fractions were collected and the STM2 and STM1 proteins detected by SDS-PAGE with Coomassie brilliant blue staining (top panels) and by immunoblotting analysis (bottom panels). Anti-Flag and Anti-Strep were used to detect Flag-STM2 and Strep-STM1, respectively.

**Extended Data Fig. 10:**
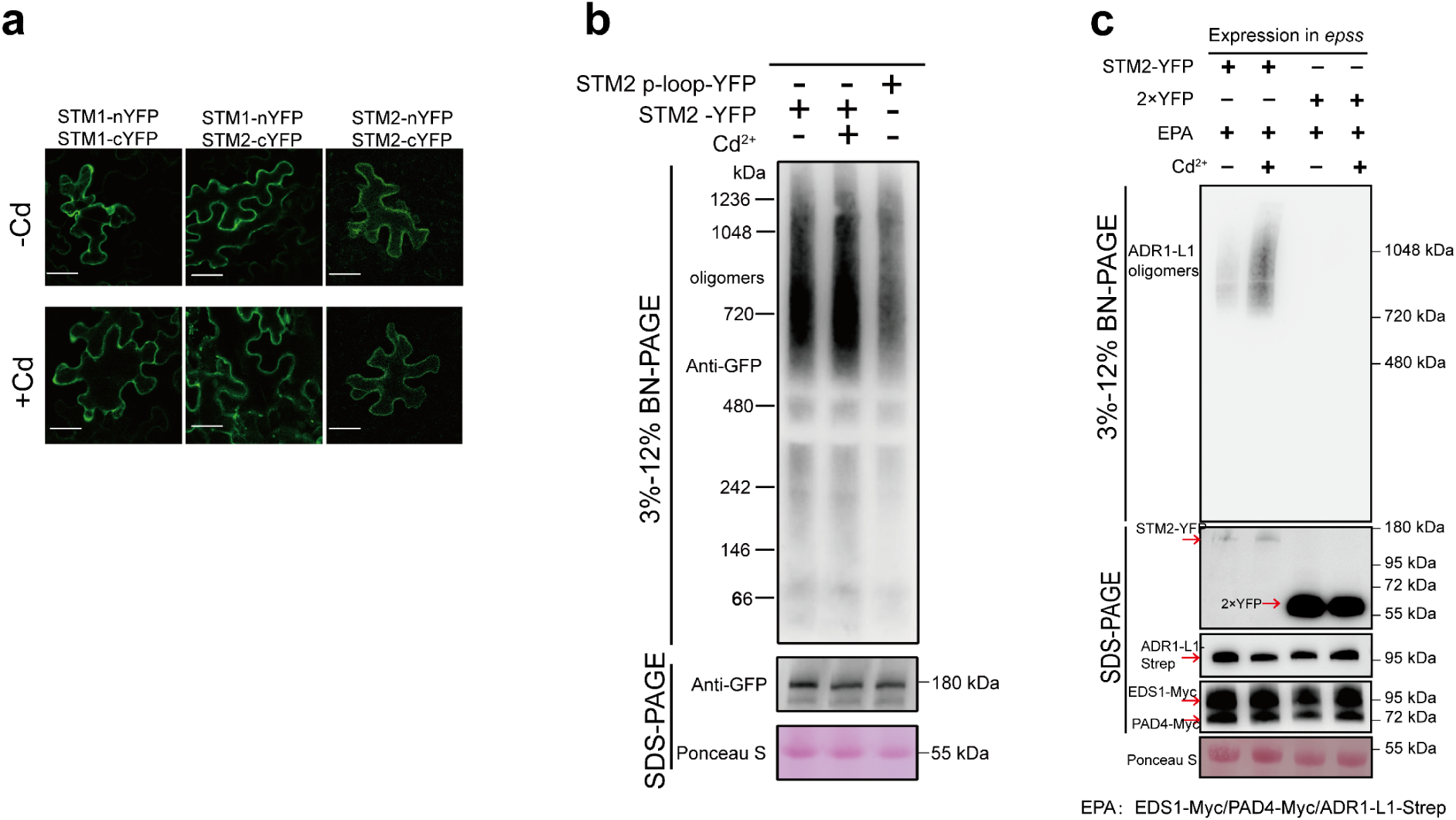
Cd promotes the production of activated ADR1-L1 oligomers via STM2. **a** BiFC assays in tobacco leaves show that Cd addition (200 µM) did not affect homomeric or heteromeric interactions between STM1 and STM2. Green fluorescence indicates the BiFC signal. Scale bar = 25 µm. **b** BN-PAGE assays show that Cd addition (200 µM) did not affect oligomerization of STM2. *STM-YFP* and the mutated p-loop motif *STM2-p-loop-YFP* (the conserved p-loop motif favors protein multimerization) constructs were transiently expressed in tobacco leaves. Cd (200 µM) or buffer solution was injected at 24 h after *Agrobacterium* injection, and samples were taken 6 h later. Protein bands between 720 and 1236 kDa indicate STM2-YFP oligomers. **c** BN-PAGE assays show that Cd promotes STM2-dependent oligomerization of AtADR1-L1. *STM2-YFP* or *YFP* was transiently co-expressed with *AtEDS1*, *AtPAD4*, and *AtADR1-L1* in tobacco *epss*. Cd (200 µM) or buffer solution was injected at 24 h after *Agrobacterium* injection, and samples were collected 6 h later. The protein bands between 720 and 1048 kDa indicate ADR1-L1-Strep oligomers.

**Extended Data Fig. 11:**
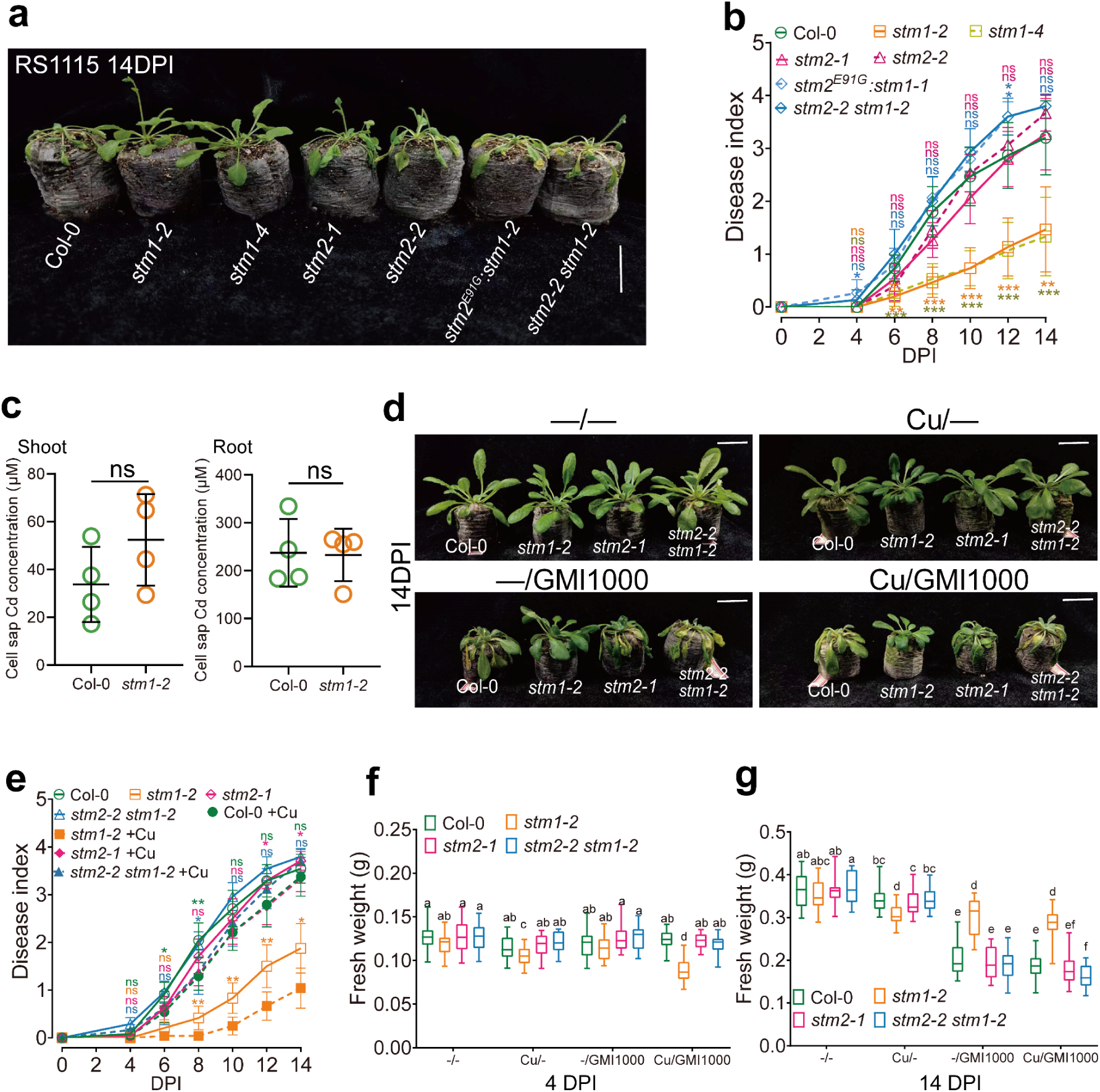
STM1 and STM2 mediate the trade-off between transition metal sensitivity and susceptibility to bacterial wilt disease. **a, b** Mutations in *STM1* enhanced resistance of *Arabidopsis thaliana* to *R. solanacearum* strain RS1115, but the resistance was lost by knockout of *STM2.* Bacterial wilt phenotype (**a**) and disease index after pathogen infection (**b**). *stm2^E91G^:stm1-2* is a single base edited line of *STM2^E91G^* in the *stm1-2* background. **c** Cd concentrations in the cell sap of shoots and roots of Col-0 and *stm1-2* plants after exposure to Cd added to the pellets for 4 days. All data points are shown with means ± SD, n=4; ns, not significant between Col-0 and *stm1-2* by two-sided *t*-test.**d-g** Interactions between *STM1/SYM2* genotype and Cu addition on the resistance to *R. solanacearum* GMI1000 infection; bacterial wilt phenotype (**d**), disease index (**e**), and plant biomass at 4 DPI (**f**) or 14 DPI (**g**). In the boxplots (**f**) and (**g**), the central line is the median, the central line represents the median, the box 25-75 percentiles and the whiskers 5-95 percentiles. Data in (**b, e**) represent means with 95% confidence interval; n = 15 in (**b**); n= 48 for 0 – 4 DPI and n = 24 for 6 – 14 DPI in (**e**). Statistical analysis was performed to compare difference from Col-0 in (**b**), and between +/- Cu in the same genotype in (**e**) by two-sided *t*-test; ns, not significant, *\* P*<0.05, ** *P*<0.01, *** *P*<0.001, and among all genotype/Cu combinations in (**f**, **g**) (3-way ANOVA, followed by Tukey’s post-hoc multiple comparisons). Different letters in (**f, g**) indicate significant difference at *P*<0.05. Scale bars = 3.5 cm in (**a, d**).

## Notes

### Competing Interest Statement

The authors have declared no competing interest.

