## Supplemental Tables 1-6 for "Transition metal-triggered immunity via an *Arabidopsis* NLR pair"

| Gene | Col-0 Control | Col-0  Cd | *stm1-2* Control | *stm1-2*  Cd | *stm2-2* Control | *stm2-2*  Cd | *stm2-2 stm1-2* Control | *stm2-2 stm1-2* Cd |
| --- | --- | --- | --- | --- | --- | --- | --- | --- |
| *PR2* | -0.76014 | 0.123617 | -0.35798 | 1.76011 | -0.76014 | -0.37019 | -0.76014 | 0.286906 |
| *ERF114* | -0.18082 | -0.42778 | -0.07561 | 1.857518 | 0.383389 | -0.39833 | -0.26135 | -0.89702 |
| *FLS2* | 0.515042 | -0.50009 | 0.144884 | 1.153343 | 0.866465 | -0.80532 | -1.10767 | -0.26665 |
| *WARKY38* | -0.01172 | -0.75292 | 1.244762 | 1.571493 | -0.65498 | -0.66539 | -0.94575 | 0.214514 |
| *AED1* | -0.24126 | -0.39976 | -0.14278 | 2.424359 | -0.41322 | -0.469 | -0.34949 | -0.40884 |
| *WARKY62* | -0.53794 | -0.70458 | 1.22308 | 2.034797 | -0.6081 | -0.44165 | -0.64959 | -0.31601 |
| *CRK6* | -0.36842 | -0.37656 | -0.30324 | 2.577042 | -0.34657 | -0.42004 | -0.36724 | -0.39497 |
| *CRK4* | 0.150716 | -1.02414 | 1.317004 | 1.518936 | 0.073709 | -0.72498 | -0.25641 | -1.05484 |
| *PNP-A* | -0.54587 | -0.45522 | -0.04151 | 2.514908 | -0.50582 | -0.3962 | -0.44732 | -0.12296 |
| *PDF1.1* | -0.4673 | -1.05743 | 0.545366 | 0.834093 | 0.13856 | -0.99325 | 0.340648 | 0.054605 |
| *PR5* | -0.61826 | -0.22924 | -0.43302 | 1.939704 | -0.42833 | -0.00159 | -0.40921 | 0.179939 |
| *SARD1* | -0.40481 | -0.77439 | 1.346899 | 1.927823 | -0.56747 | -0.52729 | -0.62632 | -0.37443 |
| *AIG1* | -0.32735 | -0.4271 | -0.14961 | 2.291697 | -0.58763 | -0.40169 | -0.53311 | -0.57459 |
| *SBT3.3* | 0.004404 | 0.290603 | -0.87337 | 1.511344 | -0.47853 | -0.16724 | -0.56845 | 0.281239 |
| *WAK1* | -0.49559 | -0.58196 | 0.997156 | 2.164765 | -0.67533 | -0.6027 | -0.6763 | -0.13005 |
| *WAK3* | -0.17748 | 0.672617 | 0.629855 | 1.56405 | -0.48454 | -0.48737 | -0.29339 | -1.42374 |
| *NIMIN1* | -0.44932 | -0.28317 | 1.091515 | 1.941913 | -0.86024 | -0.84944 | -0.63884 | 0.047583 |
| *CEP1* | -0.81179 | 0.352731 | -0.05799 | 1.708293 | -0.90846 | -0.05413 | -0.43627 | 0.20762 |
| *CRK7* | 0.322482 | -0.29243 | -0.52657 | 1.754243 | -0.52657 | -0.16302 | -0.52657 | -0.04157 |
| *CRK18* | 0.037171 | -0.54591 | -0.42457 | 1.863653 | -0.63581 | -0.6494 | -0.35833 | 0.713191 |
| *ARCK1* | -0.47253 | -0.4423 | -0.3524 | 2.263389 | -0.14212 | -0.83435 | -0.0856 | 0.065901 |
| *ABCC9* | -0.52055 | 0.246237 | 0.038964 | 2.343792 | -0.68431 | -0.52996 | -0.39979 | -0.49438 |
| *SIB1* | -0.64642 | -0.6518 | 0.101818 | 1.98523 | -0.78068 | 0.172406 | -0.58827 | 0.407719 |
| *LECRK-III.2* | -0.24784 | -0.1551 | 0.168182 | 1.829844 | 0.106308 | -0.66962 | -0.03577 | -0.996 |
| *PME12* | -0.55562 | -0.24936 | 0.153526 | 2.025702 | -1.0154 | -0.39311 | -0.33836 | 0.372615 |
| *PME12* | -0.55562 | -0.24936 | 0.153526 | 2.025702 | -1.0154 | -0.39311 | -0.33836 | 0.372615 |

**Supplemental Table 1**: Plant immunity-related genes in *stm1-2* mutant roots are significantly up-regulated by Cd exposure.

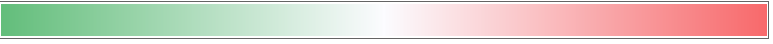

-2	0	2

FPKM Z-Score

**Supplemental Table 2**: List of plant materials used in the study.

| Name | Characteristics | Source |
| --- | --- | --- |
| *Nicotiana benthamiana* | Wild type (WT) |  |
| *epss* | *NtEDS1*, *NtPAD4*, *NtSAG101a*, *NtSAG101b* knockout *Nicotiana benthamiana.* | Jane E Parker (Max-Planck Institute) |
| Col-0 | Wild type (WT) |  |
| *cad1-3* | PCS mutant | Prof. Chris Cobbett (University of Melbourne) |
| *stm1-1 cad1-3* | Cd hypersensitive mutant for EMS mutagenic | This study |
| *stm1-2* (*SALK_004885*) | T-DNA line | Arabidopsis Biological Resource Center |
| *stm1-3 (SALK_064468)* | T-DNA line | Arabidopsis Biological Resource Center |
| *stm2-1* (*SALK_063941*) | T-DNA line | Arabidopsis Biological Resource Center |
| *rps4* (*SALK_122159*) | T-DNA line | Arabidopsis Biological Resource Center |
| *rrs1* (*SALK_131718*) | T-DNA line | Arabidopsis Biological Resource Center |
| *eds1-2* | *EDS1* mutant | Jane E. Parker (Max-Planck Institute) |
| *pad4-1* | *PAD4* mutant | Jane Glazebrook (University of Minnesota) |
| *adr1 triple* | *ADR1*/*ADR1-L1*/*ADR1-L2* triple mutant | Jeffery L. Dangl (University of North Carolina) |
| *nrg1 a/b* | *NRG1A*/*NRG1B* double mutant | Jonathan D. G. Jones(University of East Anglia) |
| *ProSTM1::STM1-GFP Com #1;Com #2；Com #3* | Transgenic complementary lines | This study |
| *AT5G45210/Com#1; Com #2* | Transgenic complementary lines | This study |
| *AT5G44370/Com#1; Com #2* | Transgenic complementary lines | This study |
| *AT5G46970/Com#1; Com #2* | Transgenic complementary lines | This study |
| *stm1-4* | CRISPR/Cas9 lines | This study |
| *stm1-5 cad1-3* | CRISPR/Cas9 lines | This study |
| *stm1-6 cad1-3* | CRISPR/Cas9 lines | This study |
| *stm1-7 cad1-3* | CRISPR/Cas9 line | This study |
| *stm2-2* | CRISPR/Cas9 line | This study |
| *stm2-2 stm1-2* | CRISPR/Cas9 lines | This study |
| *stm2-2 cad1-3* | CRISPR/Cas9 lines | This study |
| *stm2-3 stm1-2* | CRISPR/Cas9 lines | This study |
| *stm2-3 cad1-3* | CRISPR/Cas9 line | This study |
| *stm2-4* | CRISPR/Cas9 line | This study |
| *ProSTM1::GUS* | Transgenic GUS reporter line | This study |
| *ProSTM2::GUS* | Transgenic GUS reporter line | This study |
| *ProSTM1/2::GFP/mCherry::NLS* | Transgenic line | This study |
| *35S::STM1-GFP Com#1; Com#2* | Transgenic line | This study |
| *35S::STM1-GFP-NES Com#1; Com#2* | Transgenic line | This study |
| *35S::STM1-GFP-NLS Com#1; Com#2* | Transgenic line | This study |
| *STM2^E91G^:stm1-2* | single-base edited line | This study |
| *STM2^E91k^:stm1-2* | single-base edited line | This study |
| *eds1-2 stm1-2* | *eds1 stm1*double mutant from single mutant cross | This study |
| *pad4-1 stm1-2* | *pad4 stm1* double mutant from single mutant cross | This study |
| *adr1 triple stm1-2* | *adr1* *adr1-l1* *adr1-l2* *stm1* quadruple mutant from a cross between the *stm1-1* mutant and the *adr1 triple* mutant | This study |
| *nrg1 a/b stm1-2* | *nrg1a nrg1b stm1* triple mutant from a cross between the stm1-1 *mutant* and the *nrg1 a/b* mutant | This study |
| *pad4/stm1-1 cad1-3* | CRISPR/Cas9 line | This study |
| *sag101/stm1-1 cad1-3* | CRISPR/Cas9 line | This study |

**Supplemental Table 3**: Concentrations of CaCl_2_-extractable metals and pH in the soil amended with different levels of Cd.

|  | CaCl_2_-extractable metals (mg/kg) | | | | | | | |  |
| --- | --- | --- | --- | --- | --- | --- | --- | --- | --- |
| Cd addition  (mg/kg) | Cd | Mn | Fe | Co | Ni | Cu | Zn | Pb | pH |
| 0 | 0.023 | 1.550 | 0.361 | 0.039 | 0.012 | 0.206 | 1.471 | 0.014 | 5.6 |
| 1 | 0.618 | 1.297 | 0.241 | 0.027 | 0.005 | 0.160 | 0.924 | 0.006 | 5.6 |
| 2.5 | 1.357 | 1.157 | 0.203 | 0.014 | 0.007 | 0.095 | 0.943 | 0.007 | 5.6 |

**Supplemental Table 4.** List of expression constructs

| Expressed proteins | Construct | Vector | Purpose |
| --- | --- | --- | --- |
| STM1-GFP | pRCS2-STM1-GFP | initial vector: pSAT6A- EGFP- N1; final vector: pRCS2-ocs-nptII | Subcellular localization assays |
| STM2-GFP | pRCS2-STM2-GFP |  |  |
| GFP-STM1 | pRCS2-GFP-STM1 | initial vector: pSAT6A- EGFP- C1; final vector: pRCS2-ocs-nptII |  |
| GFP-STM2 | pRCS2-GFP-STM2 |  |  |
| STM1-YFP | pGWB614 -STM1-YFP | initial vector: pDONR207; final vector: pGWB614 | HR and CoIP assays |
| STM2-YFP | pGWB614 -STM2-YFP |  |  |
| Strep-STM1 | pEarleyGate-Strep-STM1 | initial vector: pDONR207; final vector: pEarleyGate |  |
| Strep-STM2 | pEarleyGate-Strep-STM2 |  |  |
| STM2-Myc | pGWB17 -STM2-Myc | initial vector: PUC19; final vector:pGWB17 | HR assays |
| RPS4-Myc | pGWB17 -RPS4-Myc |  |  |
| ROQ1-Myc | pGWB17 -ROQ1-Myc |  |  |
| Strep-stm1 | pEarleyGate-Strep-stm1 | initial vector: pDONR207; final vector: pEarleyGate | HR assays |
| EDS1-Myc | pGWB17 -EDS1-Myc | initial vector: PUC19; final vector: pGWB17 | HR and BN-PAGE assays |
| PAD4-Myc | pGWB17 -PAD4-Myc |  |  |
| ADR1-L1-Strep | pEarly 101-ADR1-L1-Strep |  |  |
| STM2E91A-YFP | pGWB614 -STM2E91A-YFP | pGWB614 | HR assays |
| STM2E91G-YFP | pGWB614 -STM2E91G-YFP |  |  |
| STM2E91K-YFP | pGWB614 -STM2E91K-YFP |  |  |
| STM2 595/599-YFP | pGWB614 -STM2 595/599-YFP |  |  |
| STM2 1022/1109-YFP | pGWB614 -STM2 1022/1109-YFP |  |  |
| STM2 Four mutant-YFP | pGWB614 - Four mutant-YFP |  |  |
| STM2 p-loop-YFP | pGWB614 - STM2 p-loop-YFP | pGWB614 | BN-PAGE |
| RPS4-TIR-Stag | pET -29a-RPS4-TIR | pET-29a | NADase in *E.coli* cells |
| STM1-TIR-Stag | pET -29a-STM1-TIR |  |  |
| STM2-TIR-Stag | pET -29a-STM2-TIR |  |  |
| STM1-nYFP | pGTQL1211YN- STM1 | initial vector: pDONR207; final vector: pGTQL1211YN | BiFC assays |
| STM2-nYFP | pGTQL1211YN- STM2 |  |  |
| STM1-cYFP | pGTQL1211YC- STM1 |  |  |
| STM2-cYFP | pGTQL1211YC- STM2 |  |  |
| Flag-STM1 | pFastBac-Flag-STM1 | pFastBac I | NADase in *vitro* and MST assay |
| Flag-STM2 | pMlink-STM2--Flag | pMlink |  |
| Flag-STM2 595/599 | pMlink-Flag-STM2-595/599 |  | MST assay |
| Flag-STM2 1022/1109 | pMlink-Flag-STM2-1022/1109 |  |  |
| Flag-STM2 Four mutant | pMlink-Flag-STM2-Four mutant |  |  |
| STM1-GFP | pFastBac- STM1-GFP | pFastBac I | Pulldown assays |
| STM1-MBP | pFastBac- STM1-MBP |  |  |
| STM2-GFP | pFastBac- STM2-GFP |  |  |
| STM2-MBP | pFastBac- STM2-MBP |  |  |

**Supplemental Table 5.** Gene accession numbers

| Gene name | Accession number | Representative transcripts |
| --- | --- | --- |
| *PCS1* | *AT5G44070* | *AT5G44070.1* |
| *STM1* | *AT5g45210* | *AT5g45210.1* |
| *STM2* | *AT5g45200* | *AT5g45200.1* |
| *EDS1* | *AT3G48090* | *AT3G48090.1* |
| *PAD4* | *AT3G52430* | *AT3G52430.1* |
| *SAG101* | *AT5G14930* | *AT5G14930.2* |
| *NRG1A* | *AT5G66900* | *AT5G66900.2* |
| *NRG1B* | *AT5G66910* | *AT5G66910.1* |
| *ADR1* | *AT1G33560* | *AT1G33560.1* |
| *ADR1-L1* | *AT4G33300* | *AT4G33300.1* |
| *ADR1-L2* | *AT5G04720* | *AT5G04720.1* |
| *PR2* | *AT3G57260* | *AT3G57260.2* |
| *PR5* | *AT1G75040* | *AT1G75040.1* |

**Supplemental Table 6.** All primers used in this study

| **Constructs for complementary test** | **Name** | **Primer (5'-3')** |
| --- | --- | --- |
| At5g45210 Gene Complementary line | gDNA-AT5G45210F: | CAGGTCGACTCTAGAGGATCCTTGGCAATCCTCTGTGAATA |
|  | gDNA-AT5G45210R: | TAAGAATTCGAGCTCGGTACCGAGTCTGATGAGCCTATGAT |
| At5g44370 Gene Complement line | gDNA-AT5G44370F: | CAGGTCGACTCTAGAGGATCCAAGTGGCTGTCACATAAGTA |
|  | gDNA-AT5G44370R: | TAAGAATTCGAGCTCGGTACCTTCACCCTCACATAATCACA |
| At5g46970 Gene Complement line | gDNA-AT5G46970F: | CAGGTCGACTCTAGAGGATCCTAACGCCCTCAATTCTCATT |
|  | gDNA-AT5G46970R: | TAAGAATTCGAGCTCGGTACCCCAGAAACAACAGCAAGAAA |
| **Constructs for tissue expression pattern and subcellular localization** | **Name** | **Primer (5'-3')** |
| *psat6a:STM1-GFP* | SAT6A-STM1 -N-F | TCAGATCTCGAGCTCAAGCTTATGGCGACTTCAAAGGCGGA |
|  | SAT6A-STM1 -N-R | CCTTGCTCACCATCAGGATCCCAGAAGATTCTTTAACATAGTATTCG |
| *psat6a:GFP-STM1* | SAT6A-STM1 -C-F | TCAGATCTCGAGCTCAAGCTTCGATGGCGACTTCAAAGGCGGA |
|  | SAT6A-STM1 -C-R | GACTCTAGACTAGGTGGATCCTTAAGAAGATTCTTTAACATAGTAT |
| *psat6a:STM2-GFP* | SAT6A-R200-N-F | TCAGATCTCGAGCTCAAGCTTATGGCGACTTCCTCCTCAAA |
|  | SAT6A-R200-N-R | CCTTGCTCACCATCAGGATCCCATCCCGCCAAAATCTAAAC |
| *psat6a:GFP-STM2* | SAT6A-R200 -C-F | TCAGATCTCGAGCTCAAGCTTCGATGGCGACTTCCTCCTCAAA |
|  | SAT6A-R200 -C-R | GACTCTAGACTAGGTGGATCCTCAATCCCGCCAAAATCTAAAC |
| *ProSTM1::GUS* | PRO-STM1-GUS-F | CTTATGCATGCGGCCGCTTAATTAATTGGCAATCCTCTGTGAATA |
|  | PRO-STM1-GUS-R | GGCGGACCTTTGCACGGCGCGCCTGATTTTCTTACTTTTTTCTTGTA |
| *ProSTM2::GUS* | PRO-STM2-GUS-F | CTTATGCATGCGGCCGCTTAATTAACTTGGAGTTCATCGTCAGTA |
|  | PRO-STM2-GUS-R: | GGCGGACCTTTGCACGGCGCGCCCAACGTTGGAGTTGTTTTTCCTT |
| *5188* | 2F:5188-GFP | ATTGTTAATTAAGAATTCGAGCTCATGGTGAGCAAGGGCGAGGAGCT |
|  | 2R：NLS+GFP: | TCATACCTTACGCTTCTTCTTTGGCTTGTACAGCTCGTCCAT |
|  | 3F:STM1UTR-NLS | CCAAAGAAGAAGCGGAAGGTATGAGCTTCCGTTCTAGGGAGGGGG |
|  | 3R: T-3STM1UTRR | ATTGTTAATTAAGAATTCGAGAGCTCAATTTTGATGGTAAAGA |
|  | 5R:5188+ RFP | AAAAACAACTCCAACGTTGTCTAGAATGGCCTCCTCCGAGAACGTC |
|  | 5F: NLS-mCherry： | TCATACCTTCCGCTTCTTCTTTGGGGGTACCAGGAACAGGTGGTGG |
|  | 6R:3STM2UTR-NLS | CCAAAGAAGAAGCGGAAGGTATGACAAGAGAAAAGGAGAGATCAAATCA |
|  | 6F:T-3UTRSTM2 | GCATGCCTGCAGGTCGACTCTTGTGGATATCTAATATTATATTA |
| *ProSTM1::STM1-GFP* | STM1-F | GGTACCGCGGGCCCGGGATCCTGATGGTGAGCAAGGGCGAGGAGCTG |
|  | GFP-NOS-R | TAAGAATTCGAGCTCGGTACCATTATGGGTATTATGGGTGC |
|  | Pro-STM1-F | CAGGTCGACTCTAGAGGATCCTGATTTTCTTACTTTTTTCTTGTA |
|  | Pro-STM1-R | CAGCTCCTCGCCCTTGCTCACCATCAGGATCCCGGGCCCGCGGTACC |
| **Constructs for changing subcellular localization** | **Name** | **Primer (5'-3')** |
| *35S::STM1-GFP* | 35S:STM1-GFP-F | TCTAGAGGATCCCCGGGTACCATGGCGACTTCAAAGGCGGA |
|  | 35S:STM1-GFP-R | GGCGGCCGCTCTAGAACTAGTTCACTTGTACAGCTCGTCC |
| *35S::STM1-GFP-NES* | 35S:STM1-GFP-F | TCTAGAGGATCCCCGGGTACCATGGCGACTTCAAAGGCGGA |
|  | gfpNES-R | TCACTTGTTAATATCAAGTCCAGCCAACTTAAGAGCAAGCTCGTTCTTGTACAGCTCGTCCATG |
|  | 35S-NES-R | GGTGGGCGGCCGCTCTAGAACTAGTTCACTTGTTAATATCAAGTCC |
| *35S::STM1-GFP-NLS* | 35S:STM1-GFP-F | TCTAGAGGATCCCCGGGTACCATGGCGACTTCAAAGGCGGA |
|  | 35S-gfpNLS-R | GTGGGCGGCCGCTCTAGAACTAGTTCATACCTTACGCTTCTTCTTTGGCTTGTACAGCTCGTCCATG |
| **Constructs for CRISPR/CAS9 knockout** | **Name** | **Primer (5'-3')** |
| Knockout of *STM1* | STM1-gRT1#U3 | TCAATGGCGACTTCAAAGGGTTTTAGAGCTAGAAAT |
|  | STM1-AtT1U3d#-: | CCTTTGAAGTCGCCATTGATGACCAATGGTGCTTTG |
|  | STM1-gRT2#U6 | CACTGACGTGGTGACGCCGGTTTTAGAGCTAGAAAT |
|  | STM1-AtT2U6-: | CGGCGTCACCACGTCAGTGCAATCACTACTTCGTCT |
| knock out of *STM2* | STM2-gRT1: | GTGAACCACTACGTTTGAGGGTTTTAGAGCTAGAAAT |
|  | STM2-AtU3RT1: | CCTCAAACGTAGTGGTTCACTGACCAATGGTGCTTTG |
|  | STM2-gRT2: | ACTTCCTCCTCAAACGTAGGTTTTAGAGCTAGAAAT |
|  | STM2-AtU6RT2: | CTACGTTTGAGGAGGAAGTCAATCACTACTTCGTCT |
| knock out of *PAD4* | PAD4-gRT1: | CAAAAGACGCGGAATGACGGGTTTTAGAGCTAGAAAT |
|  | PAD4-AtU3dT1: | CCGTCATTCCGCGTCTTTTGTGACCAATGGTGCTTTG |
|  | PAD4-gRT2: | GTCGTGGATGGAGACCACGGTTTTAGAGCTAGAAAT |
|  | PAD4-AtU6T2: | CGTGGTCTCCATCCACGACCAATCACTACTTCGTCT |
| knock out of *SAG101* | SAG101-gRT1: | CTACTACTTCTGCAGATCGGGTTTTAGAGCTAGAAAT |
|  | SAG101-AtU3bT1: | CCGATCTGCAGAAGTAGTAGTGACCAATGTTGCTCC |
|  | SAG101-gRT2: | TGAGCTATGTAAGAGACCAGGTTTTAGAGCTAGAAAT |
|  | SAG101-AtU3bT2: | CTGGTCTCTTACATAGCTCATGACCAATGTTGCTCC |
| **Constructs for single base editing** | **Name** | **Primer (5'-3')** |
| STM2 E91G | ABE-U6F: | GATTGATGAGCTGGTAAAGATCAA |
|  | ABE-U6R: | AAACTTGATCTTTACCAGCTCATC |
| STM2 E91K | CBE-U6F | GATTGCAGCTCATTTAAGCACCAT |
|  | CBE-U6R | AAACATGGTGCTTAAATGAGCTGC |
| **Constructs for transient expression** | **Name** | **Primer (5'-3')** |
| pDONR-STM1 | pDONR-STM1-F | TACAAAAAAGCAGGCTTCATGGCGACTTCAAAGGCGGATGAA |
|  | pDONR-STM1-R | GTACAAGAAAGCTGGGTCAGAAGATTCTTTAACATAGTATTCGA |
| pDONR-STM2 | pDONR-STM2-F | TACAAAAAAGCAGGCTTC ATGGCGACTTCCTCCTCAAACG |
|  | pDONR-STM2-R | GTACAAGAAAGCTGGGTCATCCCGCCAAAATCTAAA |
| pDONR-STM1-TIR | pDONR-STM1-TIR-F | GGGGACAAGTTTGTACAAAAAAGCAGGCTTCATGGCGACTTCAAAGGCGGATGAA |
| pUC19-STM1 | pUC19-STM1-F | CACCGGATCCATGGCGACTTCAAAGGCGGATGAA |
|  | pUC19-STM1-R | TTTGGATCCAGAAGATTCTTTAACATAGTATTCGA |
| pUC19-STM2 | pUC19-STM2-F | CACCGGATCCATGGCGACTTCCTCCTCAAACG |
|  | pUC19-STM2-R | TTTGAGCTCATCCCGCCAAAATCTAAA |
| pUC19-STM2 E91G | E91G-F | TGCTTAAATGGGCTGGTAAA |
|  | E91G-R | TTTACCAGCCCATTTAAGCA |
| pUC19-STM2 E91K | E91K-F | TGCTTAAATAAGCTGGTAAA |
|  | E91K-R | TTTACCAGCTTATTTAAGCA |
| pUC19-STM2 E91A | E91A-F | TGCTTAAATGCGCTGGTAAA |
|  | E91A-R | TTTACCAGCGCATTTAAGCA |
| STM2 C595A/C599A | STM2 C595A/C599A-F | TCTACAATTCTCATGCTCATAGAGAAGCTGAAGCTGAAGAC |
|  | STM2 C595A/C599A-R | GTCTTCAGCTTCAGCTTCTCTATGAGCATGAGAATTGTAGA |
| STM2 H1022 | STM2 H1022A-F | AAAACCTGCCGCGAGCTTGGAACGCAGG |
|  | STM2 H1022A-R | CCTGCGTTCCAAGCTCGCGGCAGGTTTT |
| STM2 C1109 | STM2 C1109A-F | GACGACAGTATAGGAGCTGTTGCAACCGAG |
|  | STM2 C1109A-R | CTCGGTTGCAACAGCTCCTATACTGTCGTC |
| STM2 p-loop | STM2 p-loop-F | TTGTTGGGATGCCTGCAATAGCTGAAACGGCTCTTGCCAAGAGACTAT |
|  | STM2 p-loop-R | ATAGTCTCTTGGCAAGAGCCGTTTCAGCTATTGCAGGCATCCCAACAA |
| pUC19-EDS1 | pUC19-EDS1-F | CACCGGATCCCGGCCGCCATGGCGTTTGAAGCTCTT |
|  | pUC19-EDS1-R | AAAGCTGGGTCGGCGCGCCTATCTGTTATTTCATCCATC |
| pUC19-PAD4 | pUC19-PAD4-F | CACCGGATCCCGGCCGCCATGGACGATTGTCGATTCGAGA |
|  | pUC19-PAD4-R | AAAGCTGGGTCGGCGCGCGAGTCTCCATTGCGTCACTCTCA |
| pUC19-ADR1-L1 | pUC19-ADR1-L1-F | CACCGGATCCCGGCCGCCATGGCCATCACCGATTTTTT |
|  | pUC19-ADR1-L1-R | AAAGCTGGGTCGGCGCGCGTTCGTCAAGCCAGTCTAGGCTGA |
| **Constructs for *E. coli* expression** | **Name** | **Primer (5'-3')** |
| pET29a-RPS4-TIR | 29a-RPS4-TIR-F: | GTTCCATGGCTGATATCGGATCCATGGAGACATCATCTATTTCCA |
|  | 29a- RPS4-TIR-R: | TGCTCGAGTGCGGCCGCAAGCTTTATTCCGGTCAACGCTGTCTT |
| pET29a-STM1-TIR | 29a-STM1-TIR-F: | GTTCCATGGCTGATATCGGATCCATGGCGACTTCAAAGGCGGA |
|  | 29a- STM1-TIR-R: | TGCTCGAGTGCGGCCGCAAGCTTTGTTGCATCCAGCACTTCACGT |
| pET29a-STM2-TIR | 29a-STM2-TIR-F: | GTTCCATGGCTGATATCGGATCCATGAAAGAAACCGCTGCTGC |
|  | 29a- STM2-TIR-R: | TGCTCGAGTGCGGCCGCAAGCTTTAGGATTTCCTTTACATGCTCCAC |
| **Constructs for *Sf9* expression** | **Name** | **Primer (5'-3')** |
| pFastBac-MBP | Bac1-MBP-F: | AGTCTCGAGGCATGCGGTACCCTGGTGCCACGCGGTTCCATGAAAATCGAAGAAGGTAA |
|  | Bac1-MBP-R: | CTAGTACTTCTCGACAAGCTTTCAAGTCTGCGCGTCTTTCAG |
| pFastBac-GFP | Bac1-GFP-F: | AGTCTCGAGGCATGCGGTACCCTGGTGCCACGCGGTTCCATGGTGAGCAAGGGCGAGGA |
|  | Bac1-GFP-R: | CTAGTACTTCTCGACAAGCTTTCACTTGTACAGCTCGTCCAT |
| pFastBac-STM2-GFP and pFastBac-STM2-MBP | Bac.STM2-F | CGGTCCGAAGCGCGCGGAATTCATGGCGACTTCCTCCTCAAA |
|  | Bac.STM2-R | CCGCATGCCTCGAGACTGCAGccATCCCGCCAAAATCTAAACACA |
| pFastBac-STM1-GFP and pFastBac-STM1-MBP | Bac.STM1-F | CGGTCCGAAGCGCGCGGAATTCATGGCGACTTCAAAGGCGGA |
|  | Bac.STM1-R | CCGCATGCCTCGAGACTGCAGccAGAAGATTCTTTAACATAGTAT |
| pFastBac-Flag-STM1 | FlagSTM1-F1 | CCCACCATCGGGCGCGATTACAAAGACGATGACGATAAG |
|  | FlagSTM1-F2 | GATTACAAAGACGATGACGATAAGATGGCGACTTCAAAGGCGGATG |
|  | FlagSTM1-R | TCTAGTACTTCTCGACAGAAGATTCTTTAACATAGTA |
| **Constructs for HEK293T expression** | **Name** | **Primer (5'-3')** |
| pMlink-Flag-STM2 | pMlinkSTM2-F2 | GAACAGATTGGTGGCAGTGGGATGGCGACTTCCTCCTCAAACGTA |
|  | pMlinkSUMOstar-F1 | CCTGGAAGTTCTGTTCCAGGGGCCCGACTCAGAAGTCAATCAAGAAG |
|  | pMlinkSUMOstar-R1 | TTGAGGAGGAAGTCGCCATCCCACTGCCACCAATCTGTTCTCTG |
|  | pMlinkSUMOstarSTM2-R1 | GCAAGCTTGTCGAGCCGACGCGTCCGTCACTTATCGTCATCGTCTTTGTAATC |
|  | pMlinkSUMOstarSTM2-R2 | CTTATCGTCATCGTCTTTGTAATCATCCCGCCAAAATCTAAACACA |
| **Genotyping (CRISPR/CAS9 mutants)** | **Name** | **Primer (5'-3')** |
| Identification of *STM1* knockout sites | cas9-stm1-F | TTACATTTATGGGACCCACATG |
|  | cas9-stm1-R | TACGAGTAGTTATTCTGATGTTGG |
| Identification of *STM2* knockout sites | cas9-stm2-F | CGTTTGTGTCCTTCATTTTGTACAA |
|  | cas9-stm2-R | caTTGTGACTCAGTGTACCTG |
| Identification of *PAD4* knockout sites | CAS9-PAD4-F: | GGATTGGAATTGGAATTGTTA |
|  | CAS9-PAD4-R: | GAGTTGCTGTGGTGTTGAGGA |
| Identification of *SAG101* knockout sites | CAS9-SAG-F: | TATTCCAGCTCACGCCATGGAG |
|  | CAS9-SAG-R: | GAGAGAATGATGGGTTGTTCTCGGA |
| **Genotyping (high order mutants)** | **Name** | **Primer (5'-3')** |
| Identification of *adr1* with LB_SAIL | adr1_LP : | CAAAGGCGATGATGTTCGAG |
|  | adr1_RP: | CGGATTGTTCACTATAGTAAGG |
| Identification of *adr1-11* with LB_SAIL | adr1-L1-LP: | ATGGCCATCACCGATTTTTTC |
|  | adr1-L1_RP: | GTCAGGAACAGGATTTCCAG |
| Identification of *adr1-12* with lBb1.3 | adr1-L2-LP: | ATGGCAGATATAATCGGCGG |
|  | adr1-L2-RP: | TGGGAGATTGTGACACAGTC |
| *nrg1a* mutation site | nrg1a-F: | GGAAGATGATGATGCTAGAG |
|  | nrg1a-R: | TATCAATTATTACCGAAGCACGT |
| *nrg1 b* mutation site | nrg1b-F: | CACACATCTCCAGATGAGTATG |
|  | nrg1b-R: | GGGAAGCAAGCTCATTAAGGTATAA |
| *eds1-2 mutation site* | EDS1-F | ATGGCGTTTGAAGCTCTTACCGGAA |
|  | EDS1-R | TCAGGTATCTGTTATTTCATCCATC |
| *pad4-1 mutation site* | pad4-1F | GCCAGCAAGACTCGAGATTCAATG |
|  | pad4-1R | TTCCCCTCCAAGTAATGCCCGCCA |
| **Genotyping (T-DNA mutants)** | **Name** | **Primer (5'-3')** |
| Boundary universal primers for T-DNA insertion | LB_SAIL: | TTTCATAACCAATCTCGATACAC |
|  | lBb1.3 | ATTTTGCCGATTTCGGAAC |
| Identification of *stm1-1* with LBb1.3 | *stm1-1* LP | TCTGCTTAAAGGCTGCTTCTG |
|  | *stm1-1* RP | CTCAGAGGATTACGCGTTGTC |
| Identification of *stm1-2* with LBb1.3 | *stm1-2* LP | ACGTTTGATACTCGTGCAACC |
|  | *stm1-2* RP | CTGGTGGAAATCCTAAGGCTC |
| Identification of *stm2-1* with LBb1.3 | *stm2-1* LP | ATGAGCTTGATCAAATCGTGG |
|  | *stm2-1* RP | TGTTCGCTAGCATTATCGGTC |
| Identification of *rps4* with LBb1.3 | *rps4* LP | TGCACGAACAAATAACCATTTC |
|  | *rps4* RP | TGCAAAGGAGGAAATCACATC |
| Identification of *rrs1* with LBb1.3 | *rrs1* LP | AAGTGGTGCCACAAAATCAAC |
|  | *rrs1* RP | AGAGGTCTTGAATGCACATGG |
| **Genotyping (natural variation mutants)** | **Name** | **Primer (5'-3')** |
| *cad1-3* mutation site | *pcs1*-F | TGCTGCGAACCTCTGGAA |
|  | *pcs1*-R | GATCCAGCTTTCATCCTTGC |
| The *stm1* mutation site in the *stm1-1 cad1-3* mutant | *stm1-F* | TCAGGCTACTCCACTGGGAGAAGT |
|  | *stm1-R* | CGTGCAACCAGATAGATTCAGA |
| **Primers for qPCR** | **Name** | **Primer (5'-3')** |
| Identification of expression of the endogenous gene *UBQ10-2* | QPCR-UBQ10-2 F | CGTCTTCGTGGTGGTTTCTAA |
|  | QPCR-UBQ10-2 R | GGATTATACAAGGCCCCAAAA |
| Identification of expression of the endogenous gene *ACTIN2* | QPCR-Actin2-F | TCACAGCACTTGCACCAAGC |
|  | QPCR-Actin2-R | AACGATTCCTGGACCTGCCTC |
| Identification of expression of *STM1* | QPCR-STM1-F | GTGTTGGAACTAAGAGCCTAGC |
|  | QPCR-STM1-R | TTTAGCCGATCCGTGTGTGG |
| Identification of expression of *STM2* | QPCR-STM2-F | CTATGAAAAGGGATTAGCTCTCG |
|  | QPCR-STM2-R | GAGACCACAGCACAAAGAGC |
| Identification of expression of *PR2* | QPCR-PR2-F | CGTTGTGGCTCTTTACAAACAA |
|  | QPCR-PR2-R | AGCTCTGAACGTTTTCTTGAAC |
| Identification of expression of *PR5* | QPCR-PR5-F | AGGATTTGAATTGACTCCAGGT |
|  | QPCR-PR5-R | CCATCGCCTACTAGAGTGAATT |
